# MBD2 couples DNA methylation to Transposable Elements silencing during male gametogenesis

**DOI:** 10.1101/2023.12.12.571353

**Authors:** Shuya Wang, Ming Wang, Lucia Ichino, Brandon A. Boone, Zhenhui Zhong, Ranjith K. Papareddy, Evan K. Lin, Jaewon Yun, Suhua Feng, Steven E. Jacobsen

**Affiliations:** Molecular Biology Institute, University of California Los Angeles, Los Angeles, CA 90095, USA; Department of Molecular, Cell and Developmental Biology, University of California Los Angeles, Los Angeles, CA 90095, USA; Department of Chemical and Systems Biology, Stanford University School of Medicine, Stanford, CA, USA; Eli & Edythe Broad Center of Regenerative Medicine & Stem Cell Research, University of California at Los Angeles, Los Angeles, CA 90095, USA; Howard Hughes Medical Institute, University of California Los Angeles, Los Angeles, CA 90095, USA

## Abstract

DNA methylation is an essential component of transposable element (TE) silencing, yet the mechanism by which methylation causes transcriptional repression remains poorly understood^1–5^. Here we study the *Arabidopsis thaliana* Methyl-CpG Binding Domain (MBD) proteins MBD1, MBD2, and MBD4, and show that MBD2 acts as a transposable element (TE) repressor during male gametogenesis. MBD2 bound chromatin regions containing high levels of CG methylation, and MBD2 was capable of silencing the *FWA* gene when tethered to its promoter. MBD2 loss caused TE activation in the vegetative cell (VC) of mature pollen without affecting DNA methylation levels, demonstrating that MBD2-mediated silencing acts strictly downstream of DNA methylation. Loss of silencing in *mbd2* became more significant in the *mbd5 mbd6* or *adcp1* mutant backgrounds, as well as in plants with chemically induced genome-wide DNA demethylation, suggesting that MBD2 acts redundantly with other silencing pathways to safeguard TEs from activation. Overall, our study identifies MBD2 as a novel methyl reader that acts downstream of DNA methylation to silence TEs during male gametogenesis.

---

DNA methylation at TEs usually causes transcriptional silencing, and the underlying mechanisms involve, in part, the recruitment of methyl reader proteins^1–4,6–11^. For example, in *Arabidopsis thaliana*, two functionally redundant MBD proteins, MBD5 and MBD6, bind methylated sites and prevent a subset of TEs from activation^8^. MBD1, MBD2, and MBD4 form a monophyletic group (Supplementary Fig.1a), and all three proteins contain the conserved MBD domain^8,12–14,16–18^ (Supplementary Fig.1b). While MBD2 and MBD4 possess two conserved arginine residues predicted to form hydrogen-bond and pi-cation interactions with methylated cytosines, MBD1 contains only one of these arginines^8,12–14,16–18^ (Supplementary Fig.1b).

Previous studies have concluded that MBD1, MBD2, and MBD4 lack methyl binding capacity from *in vitro* experiments^12–17^. However, post-translational modifications and buffer conditions can alter a protein’s behavior *in vitro*^19–21^, and the localization of MBD1, MBD2, and MBD4 *in vivo* has not been determined. We generated transgenic lines expressing full-length MBD1, MBD2, and MBD4 driven by their endogenous promoters and performed Chromatin Immunoprecipitation Sequencing (ChIPseq) to investigate their *in vivo* binding patterns. In line with the modeling results, MBD2 and MBD4 displayed strong enrichment at highly methylated regions, including heterochromatic TE regions and TEs associated with RNA-directed DNA methylation (RdDM) (Fig.1a, Supplementary Fig.1c-f). Furthermore, MBD2 and MBD4 exhibited a positive correlation with methylation density genome-wide (Fig.1b). In contrast, MBD1 did not enrich at methylated loci (Fig.1a-b, Supplementary Fig.1f).

**Fig 1.**
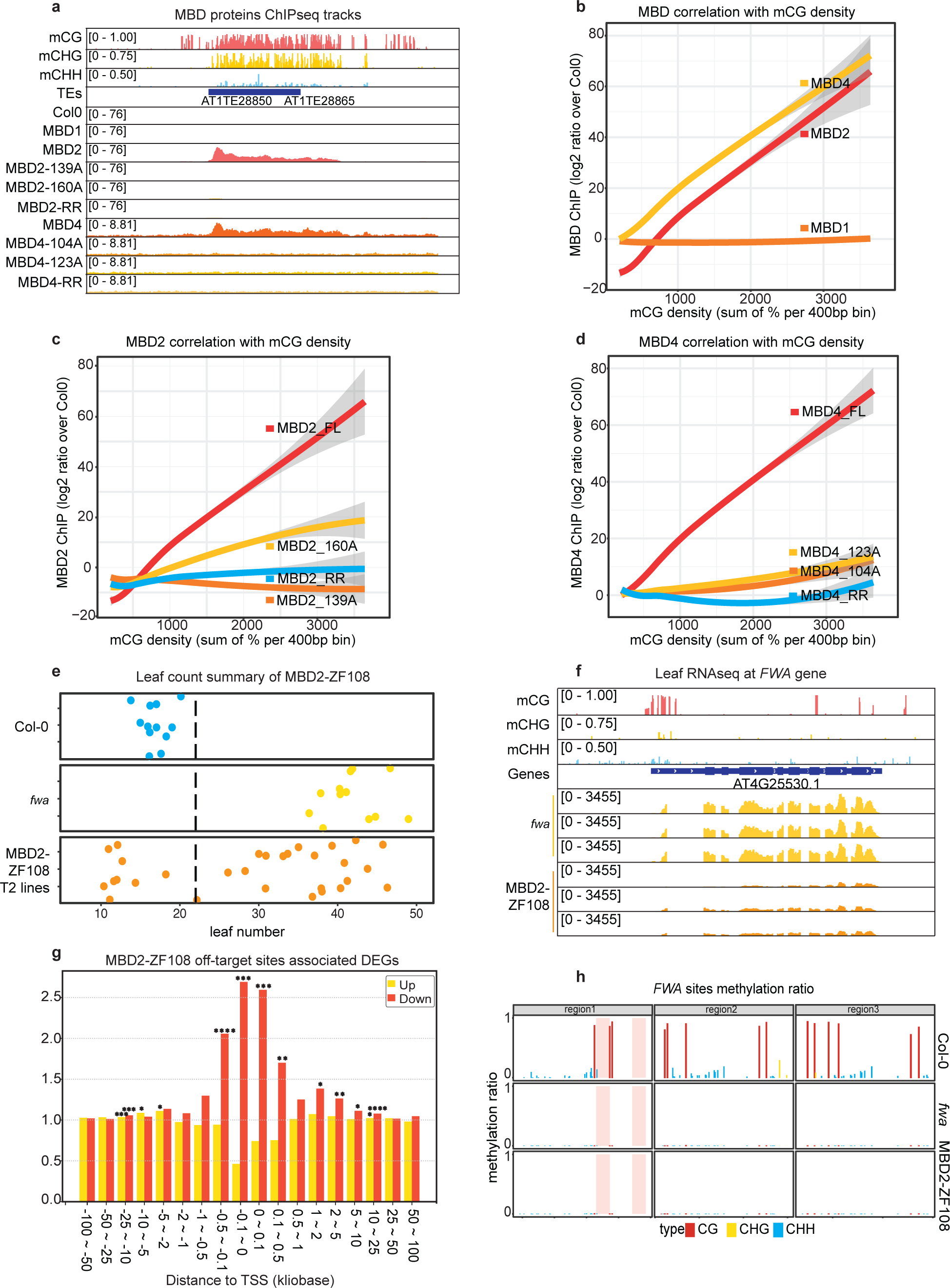
MBD2 is a novel methyl reader that silences FWA exogenously. **a.** Screenshot of the ChIP-seq tracks of MBD1, MBD2, MBD4, and the corresponding arginine mutants, together with the wild-type methylation level at this representative TE site. **b-e.** The correlation (calculated by loess regression) between CG methylation density and ChIP-seq signal of **b.** MBD1, MBD2, and MBD4. **c.** MBD2 and its arginine mutants. **d.** MBD4 and its arginine mutants. **e.** Flowering time of Col-0, *fwa*, and MBD2-ZF018 T2 lines as measured by the number of leaves. **f.** Screenshot of the leaf RNA-seq tracks of *fwa* and three representative T2 lines of MBD2-ZF108, with wild-type methylation level at the FWA locus as reference. **g.** The observed versus expected values of the upregulated (yellow bar) and downregulated (red bar) DEGs close to the ZF108 off-target sites, measured by the RAD analysis^54^. Asterisks represent the P value from the one-side hypergeometric test, P < 0.05: *; P < 0.01: **; P < 0.001: ***; P < 0.0001: **** **h.** CG, CHG, and CHH methylation at the *FWA* promoter region of Col-0, *fwa*, and a representative T2 line of MBD2-ZF108 measured by BS-PCR-seq.

To examine the necessity of the conserved arginines, we mutated either or both arginines to alanine and assessed whether MBD2 and MBD4 lost their enrichment at methylated regions. Indeed, mutations in either arginine substantially diminished the methyl binding capacity of MBD2 and MBD4, while loss of both arginines completely abolished their preference for methylated chromatin (Fig.1a, 1c-d). We also investigated the patterning of MBD2 and MBD4 at well positioned nucleosomes within heterochromatin, since the deposition of CG methylation can be guided by nucleosomes at the replication fork^1–4^. MBD2 and MBD4 peaked at the center of well-positioned nucleosomes, displaying a periodic binding pattern reflecting CG methylation density (Supplementary Fig.1g-i). In summary, these results show that MBD2 and MBD4 bind methylated CG sites in the genome, in part utilizing the conserved arginine residues thought to interact with methylated cytosines.

We next tested whether MBD1, MBD2, or MBD4 could mediate transcriptional silencing when ectopically tethered to a gene promoter. Each MBD protein was fused with an artificial zinc finger domain (ZF108) designed to bind to the target gene *FWA*^24^. This gene encodes a transcription factor normally DNA methylated at its promoter and silent in vegetative tissues.

However, Arabidopsis *fwa* epigenetic alleles have permanently lost the promoter DNA methylation, resulting in overexpression of *FWA* and a late-flowering phenotype in which plants produce an increased number of leaves prior to flowering^23^. The MBD-ZF108 fusions were transformed into *fwa*, but only the MBD2-ZF108 fusion, but not the MBD1-ZF108 or MBD4-ZF108 fusions, effectively silenced *FWA* and caused an early flowering phenotype (Fig.1e-g, Supplementary 2a-e).

ZF108 is known to bind not only the *FWA* gene but also many off-target sites genome-wide^25^. We performed Region Associated DEG (RAD) analysis^54^ at off-target sites and found that MBD2-ZF108 efficiently silences genes close to the ZF108 off-target site (Fig.1g, Supplementary 2f). To test if MBD2-ZF108-induced gene silencing was associated with changes in DNA methylation, we performed bisulfite sequencing PCR (BS-PCR) and whole-genome bisulfite sequencing (WGBS). MBD2-ZF108 did not alter the methylation status of the *FWA* promoter or of the other ZF108 binding sites, indicating that MBD2-mediated silencing is independent of DNA methylation (Fig.1h, Supplementary Fig.2h). These results show that MBD2 can act as a silencing factor downstream of DNA methylation when tethered to promoters.

Two recent studies reported that an *mbd1 mbd2 mbd4* (*mbd124)* triple mutant showed upregulation at genes involved in biotic and abiotic stress in seedlings^26–27^, but no derepression was observed at methylated TEs where methyl readers typically bind and silence^26–27^. Because these previous studies were done using vegetative tissues and because MBD5 and MBD6 have been shown to prevent TE activation specifically in pollen cells^28^, we hypothesized that MBD2 plays a similar role in pollen cells. The vegetative nucleus of pollen (VN) is known to display decompacted heterochromatin due to reduced CG methylation, H1, and H3K9me2^3–4, 28–32^ (Supplementary Fig.3a), and this partially compromised silencing creates a sensitized background in which further loss of silencing factors can cause more significant TE depression^28^. To examine whether the loss of MBD1, MBD2, and MBD4 induces transcriptional activation in pollen, and determine potential synergistic functions among the MBDs, we generated *mbd1*, *mbd14*, *mbd2*, and *mbd124* mutants and performed RNA-seq using mature pollen. We found that both CRISPR and T-DNA mutants of *mbd2* caused substantial TE reactivation (Supplementary Fig.3b). In addition, TE upregulation was rescued by re-introducing FLAG and MYC tagged MBD2 into the CRISPR mutant, demonstrating a direct role of MBD2 in repressing TEs (Supplementary Fig.3c). Consistent with ZF108 fusion experiment, only the *mbd2* and *mbd124* mutants, but not *mbd1* or *mbd14*, triggered TE expression (Fig.2a-c). Furthermore, *mbd2* and *mbd124* induced a comparable level of derepression at the same TE sites (Fig.2a-c). These results indicate that it is only MBD2 that plays a major role in preventing TE activation in mature pollen.

**Fig 2.**
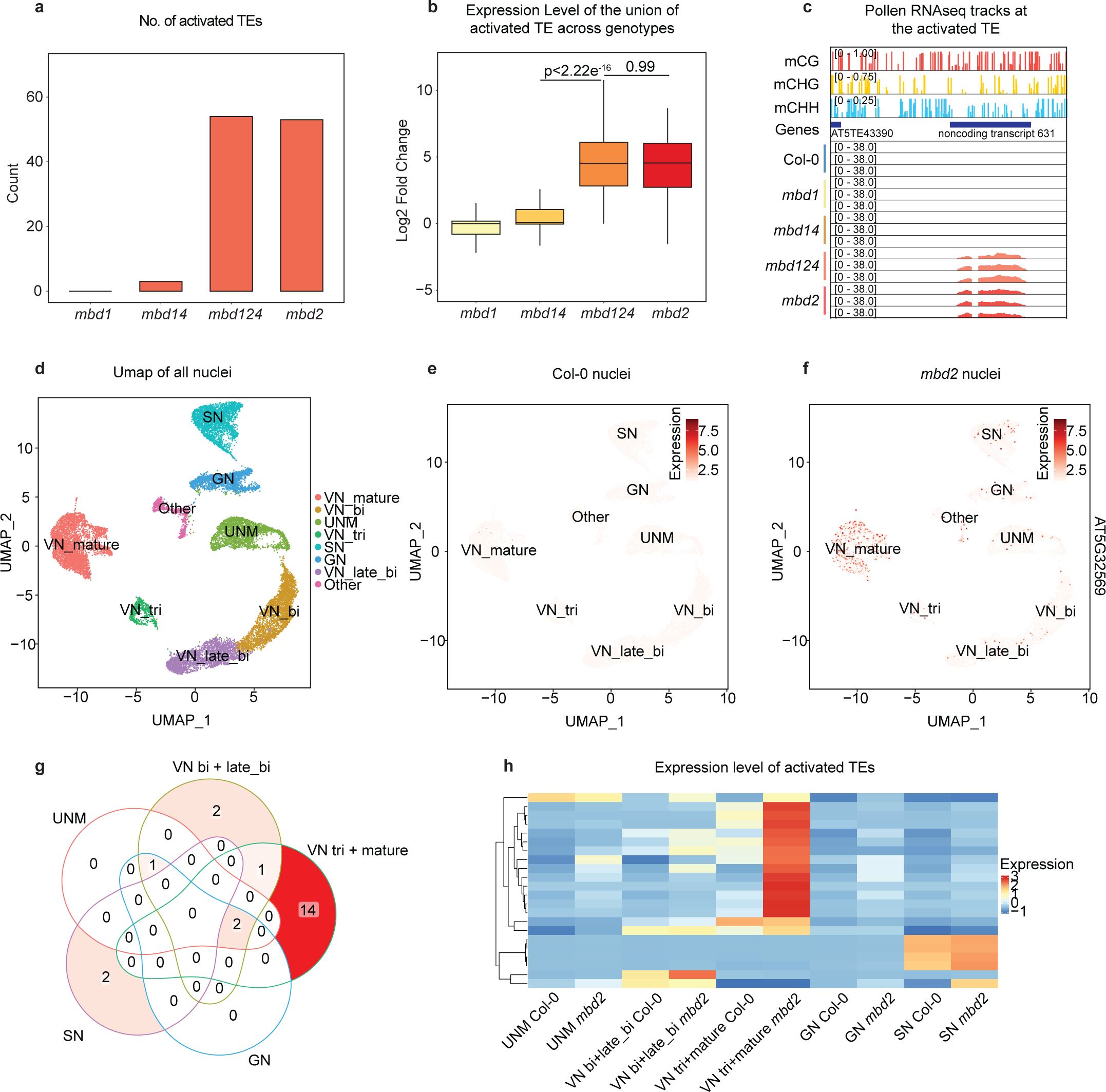
MBD2 silences TE during male gametogenesis. **a.** Count of the activated TEs from mature pollen RNA-seq of *mbd1*, *mbd14*, *mbd124*, and *mbd2* mutants. **b.** Log2 fold change of the activated TEs in *mbd1*, *mbd14*, *mbd124*, and *mbd2* mutants. P values calculated by parametric t-test are indicated. **c.** Screenshot of mature pollen RNA-seq tracks of Col-0, *mbd1*, *mbd14*, *mbd124*, and *mbd2* with wild-type methylation level at the representative TE and the methylated noncoding transcript. **d.** Umap of the integrated Col-0 and *mbd2* snRNA-seq data with cluster annotations. **e-f.** Umaps of Col-0 and *mbd2* snRNA-seq showing the expression level of the representative methylated transcript across clusters. **g.** Venn diagram showing the overlap of the activated TEs among different Umap clusters. **h.** Heatmap showing the expression of the activated TE across different clusters of Col-0 and mbd2 snRNA-seq. The expression level represents the scaled cluster averages.

To stage MBD2-mediated TE silencing during gametogenesis, we employed single-nuclei RNAseq (snRNA-seq) in wild-type Columbia-0 (Col-0) and *mbd2*. This approach allowed us to distinguish between different nuclei types involved in male gametogenesis, including microspores (UMN), generative nuclei (GN), sperm nuclei (SN), and VN^28^ (Fig.2d). We observed prominent derepression of DNA methylated TEs in tricellular and mature VN nuclei in the *mbd2* mutant (Fig.2e-h, Supplementary 3e). Interestingly, this pattern was different than that previously seen in *mbd5 mbd6* mutants, in which derepression was more prominent in the early stages of VN development^28^. Thus these different methyl readers coordinate stage-specific TE repression to safeguard the genome during the maturation of male gametophytes.

To test if the loss of MBD2 affects DNA methylation, we compared the methylation levels of mature pollen of Col-0 and the *mbd2* mutant using WGBS. We detected no genome-wide changes in DNA methylation levels, suggesting that the loss of MBD2 does not impact global DNA methylation (Supplementary Fig.3f-g). We also examined the TE sites that are activated in *mbd2*, and again found no changes in DNA methylation levels (Supplementary Fig.3h). These results indicate that MBD2 functions as a methyl reader that acts strictly downstream of DNA methylation, and is not involved in the maintenance of DNA methylation patterns.

Given that MBD2 is expressed in seedling tissues^12–16, 26–27^, and yet *mbd2* mutants showed no effect on TE silencing in this tissue^26–27^, we hypothesized that redundant silencing pathways might mask TE activation in the *mbd2* mutants in seedlings. To test this, we used the methylation inhibitor 5-azacytidine (5-AZA) to partially cripple DNA methylation in seedlings, so that we could examine the effect of *mbd2* in this sensitized background. 5-AZA reduces DNA methylation genome-wide by covalently linking to DNA methyltransferases and inhibiting their activity^33–34^. We treated Col-0 and *mbd2* seedlings with 100uM of 5-AZA and performed WGBS to examine the extent of demethylation. 100uM 5-AZA reduced DNA methylation levels by ∼30% compared to untreated controls (Fig.3a-b). We performed RNA-seq on both 5-AZA-treated and untreated Col-0 and *mbd2* seedlings, and found that more TEs are induced in *mbd2* than Col-0 in the 5-AZA treated samples (Fig.3c). Additionally, at the union of the activated TEs, *mbd2* exhibited a small increase in average TE expression (Fig.3d). These results suggest that MBD2 indeed play some roles in TE repression in seedlings, but this is only revealed when DNA methylation is partially compromised, suggesting that multiple redundant pathways are likely involved in repressing TEs in seedling tissues.

**Fig 3.**
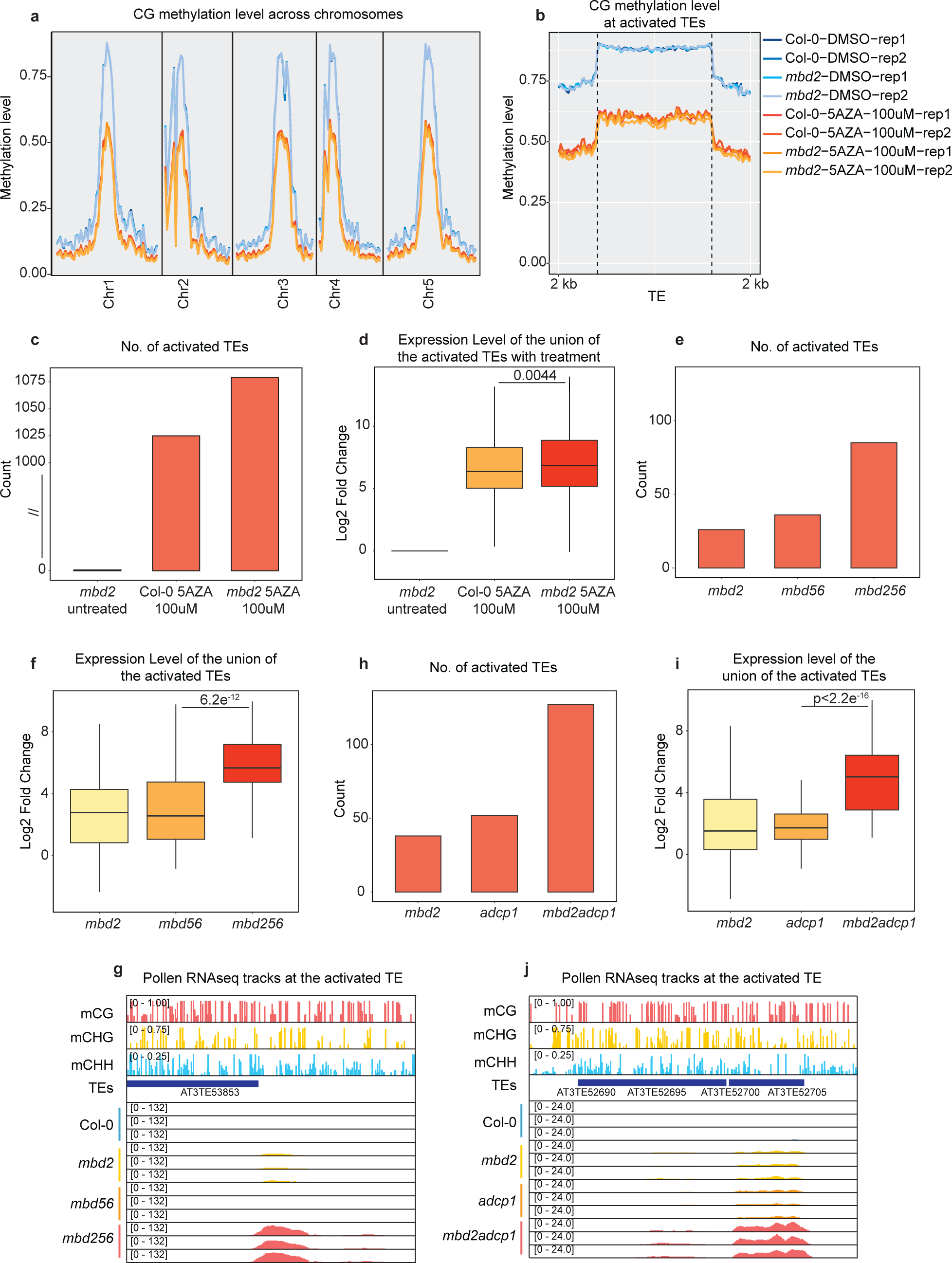
MBD2 prevents TE activation in sensitized conditions. **a.** Genome-wide CG methylation level of Col-0 and *mbd2* 10-day-old seedlings with and without 100uM 5AZA treatment. **b.** CG methylation level at the TEs activated in the *mbd2* mutant (with 100uM 5AZA treatment) across the indicated conditions**. c.** Count of the activated TEs in *mbd2*, Col-0 with 100uM 5AZA treatment, and *mbd2* with 100uM 5AZA treatment, from seedling RNA-seq. The TE count is scaled between 0 and 1000 to reflect the difference. **d.** Log2 fold change of the activated TEs in the indicated conditions. P values calculated by parametric t-test are indicated. **e.** Count of the activated TEs from mature pollen RNA-seq of mbd2, *mbd56*, and *mbd256*. **f.** Log2 fold change of the activated TEs in *mbd2*, *mbd56*, and *mbd256*. P values calculated by parametric t-test are indicated. **h.** Count of the activated TEs from mature pollen RNA-seq of *mbd2*, *adcp1*, and *mbd2 adcp1*. **i.** Log2 fold change of the activated TEs in *mbd2*, *adcp1*, and *mbd2 adcp1*. P values calculated by parametric t-test are indicated. **g-j.** Screenshots of mature pollen RNA-seq tracks of Col-0 and the indicated mutants with wild-type methylation level at the representative TE sites.

To further study possible redundancies of MBD2 silencing with other pathways, we combined *mbd2* with *mbd5 mbd6* since both MBD2 and MBD5/6 prevent TE activation during male gametogenesis. We performed mature pollen RNA-seq experiments comparing an *mbd2 mbd5 mbd6* (*mbd256*) triple mutant with *mbd2* or *mbd5 mbd6* (*mbd56*) mutants.The *mbd256* triple mutant showed derepression at a higher number of TEs than in either *mbd2* or *mbd5 mbd6* mutants (Fig.3e). Furthermore, by examining the union of TEs activated in all mutants, *mbd256* displayed an ∼8-fold higher activation relative to *mbd2* or *mbd56* mutants (Fig.3f-g). Such an enhancement was also observed in inflorescence tissues (Supplementary Fig.4a-c). These data suggest that MBD2 silences TEs redundantly with MBD5/6.

We also considered that MBD2-mediated silencing might act redundantly with silencing pathways related to the epigenetic mark H3K9me2, a hallmark of Arabidopsis heterochromatin. A previous study reported that Agenet Domain Containing Protein 1 (ADCP1), an H3K9me2 reader, maintains silencing and the integrity of the heterochromatin compartment via a liquid-liquid phase separation (LLPS) related mechanism^35^. We, therefore, generated an *mbd2 adcp1* double mutant using CRISPR-Cas9 and performed mature pollen RNA-seq of Col-0, *mbd2*, *adcp1*, and *mbd2 adcp1* mutants. We observed a dramatic enhancement in the *mbd2 adcp1* double mutant compared to the *mbd2* and *adcp1* single mutants (Fig.3h-j). The *mbd2 adcp1* double mutant induced TE derepression at ∼100 more sites than in the single mutants, and also exhibited a much higher degree of derepression at the co-activated TEs (Fig.3h-i). Moreover, we found that *mbd2 adcp1* caused activation in more TEs than did *adcp1* in inflorescence tissues, though to a lesser extent than mature pollen (Supplementary Fig.4d-f). These findings suggest that MBD2 and ADCP1 redundantly suppress a group of heterochromatic repeats, and either component is sufficient to keep these regions silent.

Previous work has suggested that HDA6 and SANT3 cooperate with MBD1, MBD2, and MBD4 to regulate protein-coding gene expression involved in stress and flowering control in seedlings, and affinity purification-mass spectrometry has demonstrated interactions between MBD2, SANT3, and HDA6^26–27^. This led us to test whether a similar mechanism could underlie MBD2-mediated TE silencing in the VN. To dissect the individual and collective roles of these proteins in TE silencing, we performed mature pollen RNA-seq on *mbd2*, *hda6*, and a *sant1 sant2 sant3 sant4* quadruple mutant (*sant1234*)^26^. Principal Component Analysis (PCA) showed that the *sant1234* samples clustered closely with the Col-0 controls suggesting that the SANT family is not involved in heterochromatin TE silencing (Fig.4a). Furthermore, *mbd2* and *hda6* samples diverged from Col-0 but formed two distinct clusters (Fig.4a), suggesting that *mbd2* and *hda6* likely induce TE activation differently. While the number of derepressed TEs in *sant1234* was very small, *mbd2* and *hda6* mutants led to activation at more than 40 TEs (Fig.4b). Among the activated TEs in *mbd2*, only around 50% were also activated in *hda6* (Fig.4c). In addition, although *hda6* led to a slightly higher degree of upregulation at the union of the activated TEs (Fig.4d, Supplementary 5a), we found that within the group of TEs activated in *mbd2*, there was a nearly 8-fold increase in TE transcript levels in *mbd2* compared to in *hda6* (Fig.4e-f, Supplementary Fig.5a). These data suggest that, while HDA6 is clearly important in TE silencing in pollen, HDA6 is likely not the primary mechanism through which MBD2 represses TEs.

**Fig 4.**
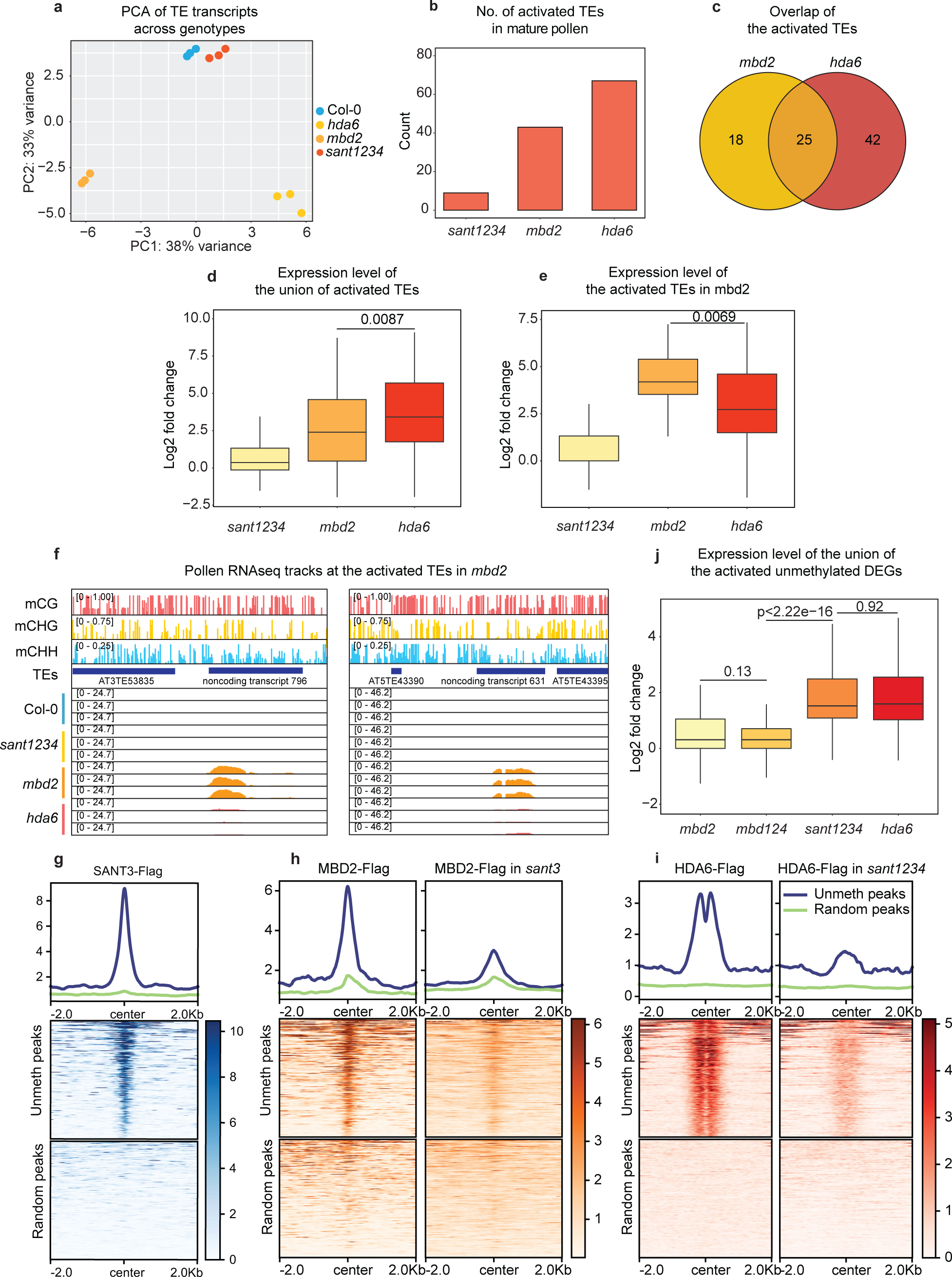
MBD2-mediated TE silencing is independent of HDA6 and SANT family proteins. **a.** PCA analysis of Col-0, *mbd2*, *sant1234*, and *hda6* based on the TE transcript from mature pollen RNA-seq. **b.** Count of the activated TEs from mature pollen RNA-seq of *mbd2*, *sant1234*, and *hda6*. **c.** Overlap of the activated TEs in *mbd2* and *hda6*. **d-e.** Log2 fold change of the **d.** union of the activated TEs **e.** mbd2 activated TEs in *mbd2*, *sant1234*, and *hda6*. P values calculated by parametric t-test are indicated. **f.** Screenshots of mature pollen RNA-seq tracks of Col-0, *mbd2*, *sant1234*, and *hda6* with wild-type methylation level at the representative TEs and methylated noncoding transcripts. **j.** Log2 fold change of the activated unmethylated DEGs in *mbd2*, *mbd124*, *sant1234*, and *hda6*. P values calculated by parametric t-test are indicated. **g-i.** Metaplots and heatmaps showing ChIP-seq signal of SANT3, MBD2, MBD2 in *sant3*, HDA6, and HDA6 in *sant1234* at the shared unmethylated peaks and random peaks.

To further examine the relationship between MBD2 and HDA6, we generated transgenic lines expressing flag-tagged HDA6 or MBD2 in either Col-0 or mutant backgrounds and performed ChIP-seq using inflorescence tissues. At genomic regions that showed ChIP-seq signals for both HDA6 and MBD2 in Col-0, and that are also DNA methylated, we observed that HDA6 ChIP-seq signals were not diminished in the absence of MBD2 and that MBD2 ChIP-seq signals were not diminished in the *hda6* mutant (Supplementary Fig.5b-d). These results suggest that MBD2 does not recruit HDA6 to its targeted regions, or vice versa. These ChIP-seq data are consistent with the RNA-seq results and suggest that HDA6 is not responsible for MBD2-mediated TE repression.

In light of the involvement of SANT family proteins and HDA6 in unmethylated gene regulation in seedlings^26–27^, we sought to dissect the formation of the SANT/HDA6/MBD2 complex and further explore its biological functions. Utilizing ChIP-seq, we identified around 1000 euchromatic peaks that did not overlap with TEs, did not contain DNA methylation, and were shared among SANT3, MBD2, and HDA6 (Fig.4g-i). Interestingly, these peaks were primarily found at the +1 nucleosome of moderately transcribed genes marked by active histone modifications, including H3K4me3 and H3K9ac (Supplementary Fig.6a-d). Additionally, HDA6 displayed a dual and symmetric peak pattern around the +1 nucleosome at these sites, which is consistent with recent structural insights on dimerized yeast HDACs bound at the two sides of a mononucleosome to access H3/H4 and H2B tails simultaneously^36^(Fig.4i). We found that the ChIP-seq signals of SANT3 and HDA6 in the *mbd2* mutant were unaltered at the ∼1000 shared regions, demonstrating that MBD2 is dispensable for the localization of SANT3 and HDA6 (Supplementary Fig.6e-f). On the other hand, in the absence of SANT3, MBD2 exhibited a dramatic reduction at these peaks (Fig.4h). Additionally, since HDA6 interacts with all SANT family proteins^26–27^, we examined HDA6 enrichment at these peaks in the *sant1234* mutant, and found that the majority of HDA6 signal was lost at these sites (Fig.4i). Collectively, these ChIP-seq results suggest that SANT3, potentially along with other SANT proteins, recruits MBD2 and HDA6 to unmethylated genes. On the other hand, the *sant3* mutant had no effect on the localization of MBD2 at its methylated target sites (Supplementary Fig.7a). Moreover, while the MBD2 double arginine mutation caused a loss of binding to DNA methylated regions, it had no effect on the localization of MBD2 to its unmethylated targets (Supplementary Fig.7b). These results highlight that MBD2 recruitment to unmethylated and methylated sites operates by different mechanisms: MBD2 binding to unmethylated sites requires SANT3 while binding to methylated sites does not involve SANT3 but requires critical amino acids in the methyl binding domain.

Since MBD2 regulates TEs in pollen, we sought to test if MBD2, together with SANTs and HDA6, might also act to repress unmethylated genes in pollen. Revisiting the mature pollen RNA-seq of Col-0, *mbd2*, *sant1234*, and *hda6* mutants, we found that while *mbd2* showed activation of only a few unmethylated genes, *sant1234*, and *hda6* showed upregulation of a much larger set (Fig.4j, Supplementary Fig.8a). Furthermore, while there was a large overlap in the upregulated genes found in *sant1234* and *hda6*, there was no overlap between these genes and genes upregulated in *mbd2* (Supplementary Fig.8a). Considering the potential redundancy with MBD1 and MBD4, we also reanalyzed the mature pollen RNA-seq of *mbd124* and again found only a few upregulated genes, and these showed no overlap with the upregulated genes in *sant1234* and *hda6* (Fig.4j, Supplementary Fig.8a). These results suggest that MBD2 plays only a minor role at unmethylated genes in pollen, while SANTs and HDA6 play a much more prominent and MBD2 independent role. This is consistent with previous work showing that the *mbd124* triple mutant caused a much weaker activation of protein-coding genes than the *sant1234* and *hda6* mutants in seedling tissues^26–27^. Overall, these results indicate that, while MBD2 is localized to some unmethylated genes, and interacts with SANT and HDA6 proteins that are also present at these genes, MBD2 has little function at these sites and mainly functions as a repressor of DNA methylated TEs.

In summary, this work demonstrates that MBD2 functions as a methyl-reader that maintains TE silencing in pollen. MBD2 silences TEs downstream of DNA methylation through a mechanism that does not require the SANT or HDA6 proteins. MBD2-mediated silencing is also distinct from the MBD5/6 and ACDP1 silencing pathways. These results highlight a high degree of redundancy between different silencing pathways acting downstream of DNA methylation, each contributing to the critical and immense function of maintaining TE repression. This multitude of silencing pathways likely reflects the intense evolutionary competition between TE proliferation and the plant genomes’ response to silence TEs and preserve genome integrity.

## Methods

### Phylogenetic Analysis

Highly conserved MBD domain sequences of MBD1, MBD2, MBD4, MBD5, MBD6, MBD7, MBD8, MBD9, MBD10, MBD11, and human MeCP2 were taken for phylogenetic analysis. All the sequences were listed in Supplementary Table S2. Protein sequence alignments were performed using ClusterOmega^56–58^. Graphic representation of the phylogenetic tree was generated using iTOL (v 6.7.5)^59^. Human MeCP2 was clustered as an outgroup given its evolutionary distance to Arabidopsis MBDs.

### Plant materials and growth conditions

The plants used in this paper were Arabidopsis thaliana Col-0 ecotype and were grown under long-day conditions (16 h light and 8 h dark). For 5AZA treatment, 100mg of 5AZA (Sigma) was first dissolved in 10ml of DMSO. Then the 5AZA stock was added to MS agar plates to reach 100uM final concentration. Control plates were only added with the same volume of DMSO.

Seedlings of the Col-0 and *mbd2* CRISPR mutant were harvested after 10-day incubation under long-day conditions. The T-DNA insertion lines used in this study are listed here: *mbd1* (SALK_025352), *mbd2* (GABI_650A05), *mbd4* (SALK_042834), *mbd6* (SALK_043927), *hda6* (SALK_201895C), and *sant3* (SALK_004966). CRIPSR mutants were generated using pYAO::hSpCas9 system^60^. *mbd2* CRISPR mutant was generated using guides: ACCGTAAATGCCCCGATAGA and CTAGGTACGCCAACCGAGTC. mbd5 CRISPR mutant was generated using guides: TCACGGAAACGTGCGACGCC and ACTTAGTATTTACTGATCGT. *adcp1* CRISPR mutant was generated using the same guides as Zhao, et al.^35^: ATTCCGCGGCTCGTGGTACATGG and GGCAGCTACCACTGAAAGGAGGG. The *sant1234* mutant is from Jian-Kang Zhu and Cui-Jun Zhang’s group^26^. Detailed information of high-order mutants generated in this study is summarized in Supplementary Table S3. Transgenic plants were generated through floral dipping using Agrobacterium (AGL0 strain).

### Plasmid construction

The Gateway-compatible binary destination vector, pEG302-effector (gDNA)-3xFLAG, was used to generate FLAG-tagged proteins for ChIP-seq experiments. The plasmid contains a Gateway cassette, a C-terminal 3xFLAG epitope tag, a Biotin Ligase Recognition Peptide, and an OCS terminator. Genomic sequences starting from the native promoter (the sequence includes ∼1.5kb upstream from the 5’ UTR or the intergenic sequences before the 5’ UTR) to the end of the endogenous gene (without stop codon) were cloned into pENTR D-TOPO vectors (Invitrogen), from which the genomic sequences were switched into the destination vector using Gateway LR Clonase II (Invitrogen). To generate constructs for ZF108 targeting experiments, pEG302-effector (gDNA)-3xFLAG-ZF108, another Gateway-compatible binary destination vector, was used. The plasmid contains a Gateway cassette, a C-terminal 3xFLAG epitope tag, a ZF108 motif, a Biotin Ligase Recognition Peptide, and an OCS terminator. The cloning strategy is the same as pEG302-effector (gDNA)-3xFLAG. pMDC123-UBQ10: effector (cDNA) ZF108-3xFLAG is also used for ZF108 targeting experiments. The plasmid, which is a Gateway-compatible binary destination vector that consists of a plant UBQ10 promoter, a C-terminal ZF108 motif, a 3xFLAG peptide, a Gateway cassette, and an OCS terminator. The cDNA was cloned first into pENTR D-TOPO vectors (Invitrogen) and then translocated into the PMDC123 destination vector via LR reaction using Gateway LR Clonase II (Invitrogen). pYAO::hSpCas9 plasmid was used to generate CRISPR mutants. Guides of MBD2, MBD5, MBD6, and ADCP1 were amplified via overlapping PCR (primer tails containing the guide sequence) using AtU6-26-sgRNA cassette as the template. Purified PCR products were cloned into pYAO::hSpCas9 plasmid via In-Fusion (Takara, 639650).

### RT-qPCR

Rosette leaf tissues from 3 to 4 week-old plants were collected. RNA was extracted using Zymo Direct-zol RNA MiniPrep kit (Zymo Research). Between 400 ng and 1 μg of total RNA was used for reverse transcription with Superscript III First Strand Synthesis Supermix (Invitrogen).

Finally, qPCR was performed with iQ SYBR Green Supermix (Bio-Rad) and FWA expression was normalized to the ISOPENTENYL PYROPHOSPHATE DIMETHYLALLYL PYROPHOSPHATE ISOMERASE 2 (IPP2). Primers are listed here:

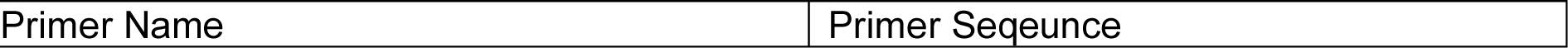
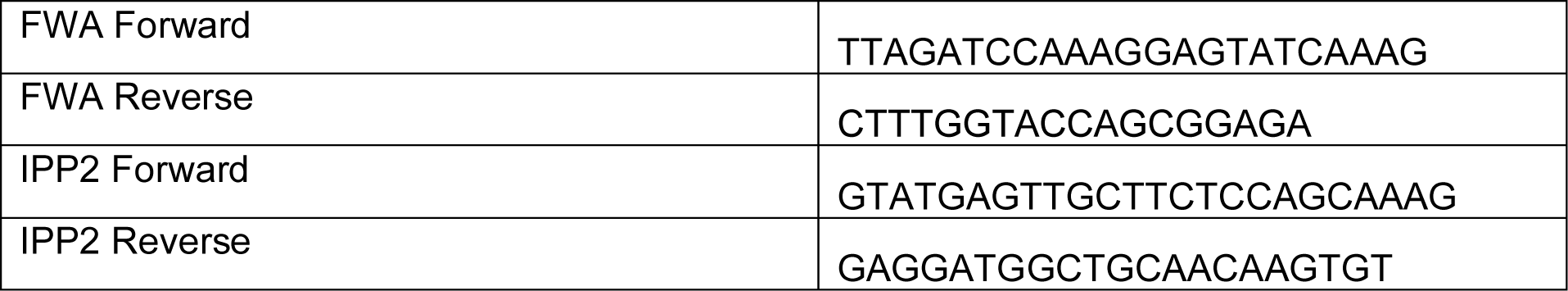

### ChIP-seq

All buffers, if not specified, were supplemented with PMSF (Sigma), Benzamindine (Sigma), and cOmpleteTM Protease Inhibitor Cocktail (Sigma). In short, 1∼2 g of unopen flower buds or 2∼4 g rosette leaves from the T2 lines were collected and ground with liquid nitrogen. 25 mL of nuclei isolation buffer (50 mM HEPES, 1 M sucrose, 5 mM KCl, 5 mM MgCl2, 0.6% Triton X-100) was used to dissolve the nuclei, and then, 680 μL of 37% formaldehyde were added to reach 1% formaldehyde concentration. The incubation lasted for 12min before adding freshly made 2M Glycine solution to quench the cross-linking. After the purification using extraction buffer 2 (0.25 M sucrose, 10 mM Tris-HCl pH 8, 10 mM MgCl2, 1% Triton X-100, 5 mM BME) and extraction buffer 3 (1.7 M sucrose, 10 mM Tris-HCl pH 8, 2 mM MgCl2, 0.15% Triton X-100, 5 mM BME), the nuclei were lysed using Nuclei lysis buffer (50 mM Tris pH 8, 10 mM EDTA, 1% SDS). The chromatin was sheared via Bioruptor Plus (Diagenode) (the setting is 30 seconds ON/30 seconds OFF, High, repeat 22 cycles). Each sample was added with 1.7 mL of ChIP Dilution Buffer (1.1% Triton X-100, 1.2 mM EDTA, 16.7 mM Tris pH 8, 167 mM NaCl) and immunoprecipitated with 10 μL of anti-FLAG (Sigma) antibody overnight at 4 °C. The next day, 50μL of Protein A and 50μL of Protein G Dynabeads (Invitrogen) were combined, washed with ChIP Dilution Buffer, and added to each sample. The incubation lasted for 2h at 4 °C. Then, 5 rounds of washes were applied to reduce the background: samples were washed twice with Low Salt Buffer (150 mM NaCl, 0.2% SDS, 0.5% Triton X-100, 2 mM EDTA, 20 mM Tris pH 8), once with High Salt Buffer (200 mM NaCl, 0.2% SDS, 0.5% Triton X-100, 2 mM EDTA, 20 mM Tris pH 8), once with LiCl Buffer (250 mM LiCl, 1% Igepal, 1% sodium deoxycholate, 1 mM EDTA, 10 mM Tris pH 8), and once with TE buffer (10 mM Tris pH 8, 1 mM EDTA). Elution was done at 65 °C using 250 μL of elution buffer (1% SDS, 10 mM EDTA, 0.1 M NaHCO3) twice.

After elution, the DNA-protein complex was reverse crosslinked using 20 μL of 5M NaCl at 65 °C overnight. The next day 1 microliter of Proteinase K (Invitrogen), 10 μL of 0.5M EDTA (Invitrogen), and 20 μL of 1M Tris pH6.5 were added to digest the proteins. Then DNA was purified using Phenol:Chloroform:Isoamyl Alcohol (Invitrogen) and precipitated with sodium acetate (Invitrogen), GlycoBlue (Invitrogen), and ethanol overnight at −20 °C. The next day, the precipitated DNA was collected and processed for the library using the Ovation Ultra Low System V2 kit (NuGEN). The sequencing was performed on Illumina NovaSeq 6000.

### MNase-seq

The buffers were supplemented with PMSF (Sigma), Benzamindine (Sigma), and cOmpleteTM Protease Inhibitor Cocktail (Sigma). Around 0.5 g of Col-0 floral tissues were harvested freshely without flash-freezing. Then 25 mL of Nuclei Isolation Buffer (300 mM Sucrose, 20 mM Tris-HCl pH 8, 5 mM MgCl2, 5 mM KCl, 0.2% Triton X-100, 5 mM BME, 35% Glycerol, 1 mM EDTA) was added and the samples were using the homogenizer (Omni International GLH) with the following setting: 1min at level 2, 45s at level 3, and 45s at level 4. Then the samples were passed through two-layer Miracloth and spinned down for 20min at 2880xg at 4 °C. After purification with extraction buffer 2 (25 M sucrose, 10 mM Tris-HCl pH 8, 10 mM MgCl2, 1% Triton X-100, 5 mM BME) and extraction buffer 3 (1.7 M sucrose, 10 mM Tris-HCl pH 8, 2 mM MgCl2, 0.15% Triton X-100, 5 mM BME), the samples were washed with 1 mL of digestion buffer (320 mM Sucrose, 50 mM Tris-HCl pH 8, 4 mM MgCl2, 1 mM CaCl2). 300 uL of digestion buffer was used to resuspend the pellet and the samples were warmed up at 37 °C for 5min.

Then 3 uL of MNase (Takara) was added to each sample and the incubation lasted for 15min at 37 °C. After digestion, 6 uL of 0.5 M EGTA (bioWORLD) and 6 uL of 0.5 M EDTA (Invitrogen) were added to quench the reaction at 65 °C for 10min. Finally 31.2 uL 5 M NaCl (Invitrogen) was used to lyse the nuclei. DNA was recovered using ChIP DNA Clean & Concentrator kit (Zymo Research) and ran on a 2% gele to check the digestion efficiency. Next the purified DNA was processed for the library using the Ovation Ultra Low System V2 kit (NuGEN). The sequencing was performed on Illumina NovaSeq 6000.

### Flowering time measurement

Rosette and cauline leaves were counted to measure the flowering time. Individual dots in the dot plots (Fig.1e, Supplementary Fig.2a-b) represent the leaf count of a single plant. The cutoff line indicates the plants with 22 or less leaves.

### Pollen extraction

Around 500 μL of open flowers were collected from 6-to 7-week-old Col-0 and the related mutants into 2.0 mL Eppendorf tubes. Then 700 μL of Galbraith Buffer (45 mM MgCl2, 30 mM C6H5Na3O7.2H2O [Trisodium citrate dihydrate], 20 mM MOPS, 0.1% [v/v] Triton X-100, pH 7) supplemented with 70mM 2-Mercaptoethanol was added, and the samples were vortexed at max speed for 3min. The suspension containing mature pollen was filtered through an 80-micron nylon mesh (Component Supply) to a new 1.5 mL Eppendorf tube. Another 700 μL of Galbraith buffer was added to the flower samples, and the above procedure was repeated. The combined 1.4 mL of samples was centrifuged at 800xg at 4 °Cand the pollen pellet was collected. A metal bead was added to each sample for later grinding. The samples were flash-frozen in liquid nitrogen.

### RNA-seq

Biological triplicates were used for each genotype. Mature pollen was harvested using the above protocol. One inflorescence containing unopen flower buds from a 5-to 6-week-old plant was collected as a biological replicate and frozen in liquid nitrogen. Rosette leaf tissues from 4-to 5-week-old plants were harvested from a single plant as biological replicate and flash-frozen in liquid nitrogen. The samples were ground into powder, and RNA was extracted using the Direct-zol RNA MiniPrep kit (Zymo Research). 250ng of total RNA from mature pollen and 1000 nanograms of total RNA from inflorescences or leaf tissues were used for RNA-seq library preparation with TruSeq Stranded mRNA kit (Illumina). The final library was sequenced on Illumina NovaSeq 6000 or HiSeq 4000 instruments.

### BS-PCR-seq

Rosette leaf tissues from 4-to 5-week-old Col-0 and representative T2 MBD2-ZF108 transgenic lines that display early flowering phenotype were collected for BS-PCR at *FWA* promoter regions. DNA was extracted using the DNeasy Plant Mini kit (Qiagen). Around 2 micrograms of DNA was used for DNA bisulfate conversion by EpiTect Bisulfate kit (QIAGEN). The converted DNA served as a template for amplification. PCR was performed at three different regions spanning over the promoter and the 5′ transcribed regions of *FWA*, including region 1 (chr4: 13038143-13038272), region 2 (chr4: 13038356-13038499), and region 3 (chr4: 13038568-13038695). The PCR reactions used Pfu Turbo Cx (Agilent), dNTP (Takara Bio), and the primers designed for the above-mentioned *FWA* regions. PCR products from the same sample were pooled and cleaned using AMPure beads (Beckman CoμLter). The purified PCR products were prepared for the libraries using the Kapa DNA Hyper Kit (Roche) utilizing indexes from TruSeq DNA UD indexes for Illumina (Illumina). Finally, the libraries were sequenced on Illumina iSeq 100.

### Single-nuclei RNA-seq

The experiment was performed following the published protocol^28^. Each buffer was freshly added with 2-Mercaptoethanl to reach 70mM concentration and supplemented with cOmpleteTM Protease Inhibitor Cocktail (Sigma). In brief, 5 mL of unopen flower buds and open flowers from the same inflorescence were harvested on ice. Prechilled mortar and pestle were used to release the spores from the buds with 5 mL of 0.1M mannitol. The liquid containing the released spores was transferred to a 50 mL conical tube. Another 10 mL of 0.1M mannitol was used to rinse the mortar and pestle. Then the samples were vortexed at max speed for 30 s to further release the spores. Then the samples were filtered through a 100-micron nylon mesh (Component Supply) to remove the debris. Another 5 mL was used to rinse the tubes and filtered through the same 100-micron nylon mesh. 20 mL of the suspension containing the spores was filtered again through a 60-micron nylon mesh. The sample was distributed into two 15 mL glass tubes and centrifuged with a Sorvall Lynx 4000 Centrifuge (Thermo Scientific) with TH13–6×50 swing-out rotor for 10min at 900xg at 4 °C. Each pellet was resuspended in 1 mL of ice-cold 0.1 M mannitol and transferred into a new tube to layer over 3 mL of 20% Percoll. The samples were centrifuged again for 10min at 450xg at 4 °C. Then the pellets from the same genotype were combined with 2 mL of 0.1M mannitol and centrifuged over 20% Percoll with the same setting for another two times. The purified mixed spores were transferred into a 1.5 mL Eppendorf tube and centrifuged for 5 min at 500xg at 4 °C. 800 μL of Galbraith buffer was used to resuspend the pellet. The samples were then transferred to a 1.5 mL tube with 100 mL of acid-washed 0.5 mm glass beads (Sigma). To break the pollen cell walls, the samples were vortexed at max speed for 2 min at 4 °C with the setting below: for the 1^st^ minute, 7 s vortex, 3 s inversion; for the 2^nd^ minute, 7 s vortex, 2 s inversion. A 10 mm cellTrics filter was placed in a clean 1.5 mL tube, and the suspension was added on and filtered by brief centrifugation. This flowthrough was kept on ice. The beads were rinsed again with 400 μL of Galbraith buffer, and the suspension was transferred to the cellTrics filter, and centrifuged to collect more nuclei.

Given that unbroken pollen grains remained on the filter, 800 μL of Galbraith buffer was added to transfer the suspension on the filter back into the tube with the glass beads. The vortexing and filtering were repeated. The combined suspension was then centrifuged for 5 min at 500 g at 4 °C, and the nuclei pellets were resuspended with 50 μL of CyStain UV Precise P - Nuclei Extraction Buffer (Sysmex, 05-5002-P02). 400 μL of CyStain UV Precise P - Staining Buffer (Sysmex, 05-5002-P01) was added to stain the nuclei, and the samples were added with Protector RNase Inhibitor (Sigma) to reach a final concentration of 0.2 U/mL. The samples were passed into a FACS tube (Falcon 352235) and sorted immediately. Sorting was performed with a BD FACS ARIAII instrument equipped with a 355nm UV laser, using the 70 mm nozzle. For each sample, 40,000–60,000 nuclei were sorted in 500 mL of Nuclei wash buffer (2% BSA in 1X PBS) supplemented with Protector RNase Inhibitor (Sigma). The sorted nuclei were centrifuged for 5 min at 500 g at 4 °C. Finally, the pellet was resuspended in 20–25 μL of buffer and sent as input for the 10x Genomics Chromium Single Cell 30 Reagent Kit v3.

### Whole-genome bisulfite sequencing

Rosette leaf tissues from 4-to 5-week-old Col-0 and representative T2 MBD2-ZF108 transgenic lines that display early flowering phenotype were collected and frozen in liquid nitrogen. For mature pollen WGBS, 1000 μL of open flowers were collected, and the pollen pellets were immediately frozen after purification. DNA from leaf tissues and mature pollen were extracted using the DNeasy Plant Mini kit (Qiagen). A total of 500ng DNA from leaf tissues and 100ng DNA from mature pollen was sheared to ∼300bp using the Covaris S2 (Covaris). Then the libraries were constructed using the Ovation Ultralow Methyl-seq kit (NuGEN), and bisulfite conversion was achieved using the Epitect Bisulfite Conversion kit (QIAGEN). Finally, the libraries were sequenced on Illumina NovaSeq 6000 instruments or HiSeq 4000 instruments.

### ChIP-seq analysis

Quality control was initially run to filter out the low-quality reads. Trim Galore (Babraham Institute) was used to remove the Illumina adapters. Then the reads were aligned to the Arabidopsis reference genome (TAIR10) using the bowtie2^46^, allowing only uniquely mapped reads with perfect matches. MarkDuplicates.jar (picard-tools suite, Broad Institute) was used to remove the PCR duplicates. Bigwig files were generated using deeptools (v 3.0.2) bamCoverage^47^ with the options -- normalizeUsing RPGC and --binSize 10. For correlation analysis between the ChIP-seq signal and mCG density, the samples were normalized to the no-FLAG control using bamCompare^47^ with the options --scaleFactorsMethod readCount, -- binSize 10, and --operation log2. The normalized ChIP-seq signal and CG methylation percentages were summarized into 400 bp bins. We took a random subset covering 10% of all genomic regions for the correlation analysis. The data were plotted using the R package *ggplot* with the option *geom_smooth*. ChIP-seq peaks were called using MACS2 (v2.1.1)^48^ using an FDR cutoff of 0.05. The FLAG-associated hyperchipable regions, defined as peaks called in the anti-FLAG Col-0 controls, were removed from the peak files. Heterochromatin peaks were defined as peaks intersecting with TAIR10 pericentromeric regions using bedtools intersect^55^ function from deeptools (v 3.0.2).

### MNase-seq analysis

The reads of low quality were fileterd out and the adaptors were trimmed with Trim Galore (Babraham Institute). Next the processed reads were aligned to TAIR10 using bowtie2^46^ keeping reads smaller than 2000 bp and allowing only uniquely mapped reads with perfect matches. Then PCR duplicates were removes using MarkDuplicate (picard-tools suit, Broad Institute) and bigwig files were generated using bamCoverage^47^.

### RNA-seq analysis

RNAs-seq reads were filtered according to quality score and were trimmed out Illumina adaptors using Trim Galore (Babraham Institute). Then the filtered reads were mapped to the Arabidopsis reference genome (TAIR10) using STAR^49^. We allowed only uniquely mapped reads with less than 5% of mismatches. Bigwig files for genome browser visualization were generated using bamCoverage^47^ with the options -- normalizeUsing RPGC and --binSize 10. HTSeq^50^ was used to obtain the read counts for TE and our previously reannotated pollen transcripts^28^. DESeq2 was used to perform the differential analysis with the cutoff padj < 0.05 and |log2FC| > 1. We used *ggplot* to generate all the related plots.

### Single-nuclei RNA-seq analysis

The analysis was performed following the published pipeline^28^. In brief, Cell Ranger 6.1.1 was used to process the raw data following the published pollen transcriptome reannotations^28^. With Cell Ranger results, SoupX^43^ and Seurat^44^ were used to remove the ambient RNA and filter out the cells detected with less than 200 genes. The data were normalized and scaled following the published settings^28^. After the normalization, PCA analysis was performed (npc=20).

DoubletFinder v3^45^ was used to identify doublets and *find.pK* was used to obtain the ideal pK parameters for each sample. The percentage of doublets removed and the pK values were summarized in Table S1. Col-0 and mbd2 datasets were integrated with Seurat v4 *FindIntegrationAnchors* and *IntegrateData* using default settings. The data was scaled, and PCA analysis was performed (npcs=40). Then clustering analysis was done using *FindNeighbors* and *FindClusters* functions. The number of cells per cluster is summarized in Table S1. In addition, the markers for each cluster were obtained with *FindAllMarker* using the integrated dataset. Finally, DEG analysis was performed on individual clusters. We specifically focused on activated TEs using the cutoff padj < 0.05 & |avg_log2FC| > 0.25. In this analysis, the following clusters were groups: VN_bi and VN_late_bi, VN_tri and VAN_mature. The TE expression heatmap was generated using Seurat function *AverageExpression*.

### Whole-genome bisulfite sequencing analysis

WGBS were filtered and removed with Illumina adaptors using Trim Galore (Babraham Institute). Reads with three or more consecutively methylated CHH sites were considered as non-converted reads and removed from the analyses. Bismark^51^ was used to map the reads to the Arabidopsis reference genome (TAIR10) and obtain the methylation percentages for each cytosine. We used ViewBS^52^ to generate the plots showing the genome-wide methylation information across genotypes.

## Data availability

The high-throughput sequencing data generated in this paper have been deposited in the Gene Expression Omnibus (GEO) database.

## Acknowledgments

We thank Gabriella Rubert, Priyanka Vaidya, Suzanne Sajid, Hanyi Jia, and Alexander Barinsky for their technical support; Dr. Yan Xue, Dr. Jason Gardiner, Dr. Colette Picard, Dr. Zheng Li, and Dr. Zhongshou Wu for discussion and advice. We also thank Mahnaz Akhavan and the UCLA BSCRC BioSequencing Core for the sequencing support. This work was supported by a George G. & Betsy H. Laties Graduate Fellowship in Molecular Plant Biology to S.W., a Philip Whitcome Pre-Doctoral Fellowship in Molecular Biology to L.I., (Brandon’s funding) and (Steve’s funding) to S.E.J. S.E.J. is Howard Hughes Medical Institute Investigator.

## Authors Contributions

S.W. and S.E.J. conceived the study, designed the research, and wrote the manuscript. S.W. performed most of the experiments and data analysis. M.W., L.I., B.A.B., R.K.P., E.K.L., and J.Y. contributed to the experiments. Z.Z contributed to the data analysis. S.F. performed BS-PCT-seq and all high-throughput sequencing.

## Reference

1. Greenberg, Maxim V. C., and Deborah Bourc’his. “The Diverse Roles of DNA Methylation in Mammalian Development and Disease.” Nature Reviews Molecular Cell Biology 20, no. 10 (October 2019): 590–607. 10.1038/s41580-019-0159-6.

2. Smith, Zachary D., and Alexander Meissner. “DNA Methylation: Roles in Mammalian Development.” Nature Reviews Genetics 14, no. 3 (March 2013): 204–20. 10.1038/nrg3354.

3. Law, Julie A., and Steven E. Jacobsen. “Establishing, Maintaining and Modifying DNA Methylation Patterns in Plants and Animals.” Nature Reviews Genetics 11, no. 3 (March 2010): 204–20. 10.1038/nrg2719.

4. Zhang, Huiming, Zhaobo Lang, and Jian-Kang Zhu. “Dynamics and Function of DNA Methylation in Plants.” Nature Reviews Molecular Cell Biology 19, no. 8 (August 2018): 489–506. 10.1038/s41580-018-0016-z.

5. Kankel, Mark W, Douglas E Ramsey, Trevor L Stokes, Susan K Flowers, Jeremy R Haag, Jeffrey A Jeddeloh, Nicole C Riddle, Michelle L Verbsky, and Eric J Richards. “Arabidopsis *MET1* Cytosine Methyltransferase Mutants.” Genetics 163, no. 3 (March 1, 2003): 1109–22. 10.1093/genetics/163.3.1109.

6. Jones, Peter L., Gert Jan C. Veenstra, Paul A. Wade, Danielle Vermaak, Stefan U. Kass, Nicoletta Landsberger, John Strouboulis, and Alan P. Wolffe. “Methylated DNA and MeCP2 Recruit Histone Deacetylase to Repress Transcription.” Nature Genetics 19, no. 2 (June 1998): 187–91. 10.1038/561.

7. Nan, Xinsheng, Huck-Hui Ng, Colin A. Johnson, Carol D. Laherty, Bryan M. Turner, Robert N. Eisenman, and Adrian Bird. “Transcriptional Repression by the Methyl-CpG-Binding Protein MeCP2 Involves a Histone Deacetylase Complex.” Nature 393, no. 6683 (May 1998): 386–89. 10.1038/30764.

8. Ichino, Lucia, Brandon A. Boone, Luke Strauskulage, C. Jake Harris, Gundeep Kaur, Matthew A. Gladstone, Maverick Tan, et al. “MBD5 and MBD6 Couple DNA Methylation to Gene Silencing through the J-Domain Protein SILENZIO.” Science 372, no. 6549 (June 25, 2021): 1434–39. 10.1126/science.abg6130.

9. Lai, Anne Y., and Paul A. Wade. “Cancer Biology and NuRD: A Multifaceted Chromatin Remodelling Complex.” Nature Reviews Cancer 11, no. 8 (August 2011): 588–96. 10.1038/nrc3091.

10. Denslow, S A, and P A Wade. “The Human Mi-2/NuRD Complex and Gene Regulation.” Oncogene 26, no. 37 (August 13, 2007): 5433–38. 10.1038/sj.onc.1210611.

11. Allen, Hillary F., Paul A. Wade, and Tatiana G. Kutateladze. “The NuRD Architecture.” Cellular and Molecular Life Sciences 70, no. 19 (October 2013): 3513–24. 10.1007/s00018-012-1256-2.

12. Zemach, Assaf, and Gideon Grafi. “Characterization of *Arabidopsis Thaliana* Methyl-CpG-Binding Domain (MBD) Proteins: *Methyl-CpG-Binding Domain (MBD) Proteins in* Arabidopsis.” The Plant Journal 34, no. 5 (June 2003): 565–72. 10.1046/j.1365-313X.2003.01756.x.

13. Scebba, Francesca, Giovanni Bernacchia, Morena De Bastiani, Monica Evangelista, Rita Maria Cantoni, Rino Cella, Maria Tereasa Locci, and Letizia Pitto. “Arabidopsis MBD Proteins Show Different Binding Specificities and Nuclear Localization.” Plant Molecular Biology 53, no. 5 (November 2003): 755–71. 10.1023/B:PLAN.0000019118.56822.a9.

14. Ito, Mikako, Akiko Koike, Nozomu Koizumi, and Hiroshi Sano. “Methylated DNA-Binding Proteins from Arabidopsis.” Plant Physiology 133, no. 4 (December 1, 2003): 1747–54. 10.1104/pp.103.026708.

15. Zemach, Assaf, Yan Li, Bess Wayburn, Hagit Ben-Meir, Vladimir Kiss, Yigal Avivi, Vyacheslav Kalchenko, Steven E. Jacobsen, and Gideon Grafi. “DDM1 Binds Arabidopsis Methyl-CpG Binding Domain Proteins and Affects Their Subnuclear Localization.” The Plant Cell 17, no. 5 (April 29, 2005): 1549–58. 10.1105/tpc.105.031567

16. Zemach, Assaf, and Gideon Grafi. “Methyl-CpG-Binding Domain Proteins in Plants: Interpreters of DNA Methylation.” Trends in Plant Science 12, no. 2 (February 2007): 80–85. 10.1016/j.tplants.2006.12.004.

17. Wu, Zhibin, Sizhuo Chen, Mengqi Zhou, Lingbo Jia, Zhenhua Li, Xiyou Zhang, Jinrong Min, and Ke Liu. “Family-Wide Characterization of Methylated DNA Binding Ability of Arabidopsis MBDs.” Journal of Molecular Biology 434, no. 2 (January 2022): 167404. 10.1016/j.jmb.2021.167404.

18. Mahana, Yutaka, Izuru Ohki, Erik Walinda, Daichi Morimoto, Kenji Sugase, and Masahiro Shirakawa. “Structural Insights into Methylated DNA Recognition by the Methyl-CpG Binding Domain of MBD6 from *Arabidopsis Thaliana*.” ACS Omega 7, no. 4 (February 1, 2022): 3212–21. 10.1021/acsomega.1c04917.

19. Macek, Boris, Karl Forchhammer, Julie Hardouin, Eilika Weber-Ban, Christophe Grangeasse, and Ivan Mijakovic. “Protein Post-Translational Modifications in Bacteria.” Nature Reviews Microbiology 17, no. 11 (November 2019): 651–64. 10.1038/s41579-019-0243-0.

20. Ytterberg, A. Jimmy, and Ole N. Jensen. “Modification-Specific Proteomics in Plant Biology.” Journal of Proteomics 73, no. 11 (October 2010): 2249–66. 10.1016/j.jprot.2010.06.002.

21. Zhang, Min, Jun-Yu Xu, Hao Hu, Bang-Ce Ye, and Minjia Tan. “Systematic Proteomic Analysis of Protein Methylation in Prokaryotes and Eukaryotes Revealed Distinct Substrate Specificity.” PROTEOMICS 18, no. 1 (January 2018): 1700300. 10.1002/pmic.201700300.

22. Stoddard, Caitlin I., Suhua Feng, Melody G. Campbell, Wanlu Liu, Haifeng Wang, Xuehua Zhong, Yana Bernatavichute, Yifan Cheng, Steven E. Jacobsen, and Geeta J. Narlikar. “A Nucleosome Bridging Mechanism for Activation of a Maintenance DNA Methyltransferase.” Molecular Cell 73, no. 1 (January 2019): 73–83.e6. 10.1016/j.molcel.2018.10.006.

23. Soppe, Wim J.J, Steven E Jacobsen, Carlos Alonso-Blanco, James P Jackson, Tetsuji Kakutani, Maarten Koornneef, and Anton J.M Peeters. “The Late Flowering Phenotype of Fwa Mutants Is Caused by Gain-of-Function Epigenetic Alleles of a Homeodomain Gene.” Molecular Cell 6, no. 4 (October 2000): 791–802. 10.1016/S1097-2765(05)00090-0.

24. Johnson, Lianna M., Jiamu Du, Christopher J. Hale, Sylvain Bischof, Suhua Feng, Ramakrishna K. Chodavarapu, Xuehua Zhong, et al. “SRA-and SET-Domain-Containing Proteins Link RNA Polymerase V Occupancy to DNA Methylation.” Nature 507, no. 7490 (March 6, 2014): 124–28. 10.1038/nature12931.

25. Gallego-Bartolomé, Javier, Wanlu Liu, Peggy Hsuanyu Kuo, Suhua Feng, Basudev Ghoshal, Jason Gardiner, Jenny Miao-Chi Zhao, Soo Young Park, Joanne Chory, and Steven E. Jacobsen. “Co-Targeting RNA Polymerases IV and V Promotes Efficient De Novo DNA Methylation in Arabidopsis.” Cell 176, no. 5 (February 2019): 1068–1082.e19. 10.1016/j.cell.2019.01.029.

26. Zhou, Xishi, Junna He, Christos N. Velanis, Yiwang Zhu, Yuhan He, Kai Tang, Mingku Zhu, et al. “A Domesticated *Harbinger* Transposase Forms a Complex with HDA6 and Promotes Histone H3 Deacetylation at Genes but Not TEs in *Arabidopsis*.” Journal of Integrative Plant Biology 63, no. 8 (August 2021): 1462–74. 10.1111/jipb.13108.

27. Feng, Chao, Xue-Wei Cai, Yin-Na Su, Lin Li, She Chen, and Xin-Jian He. “Arabidopsis RPD3-like Histone Deacetylases Form Multiple Complexes Involved in Stress Response.” Journal of Genetics and Genomics 48, no. 5 (May 2021): 369–83. 10.1016/j.jgg.2021.04.004.

28. Ichino, Lucia, Colette L. Picard, Jaewon Yun, Meera Chotai, Shuya Wang, Evan K. Lin, Ranjith K. Papareddy, Yan Xue, and Steven E. Jacobsen. “Single-Nucleus RNA-Seq Reveals That MBD5, MBD6, and SILENZIO Maintain Silencing in the Vegetative Cell of Developing Pollen.” Cell Reports 41, no. 8 (November 2022): 111699. 10.1016/j.celrep.2022.111699.

29. Borg, M., L. Brownfield, and D. Twell. “Male Gametophyte Development: A Molecular Perspective.” Journal of Experimental Botany 60, no. 5 (January 23, 2009): 1465–78. 10.1093/jxb/ern355.

30. Berger, Frédéric, and David Twell. “Germline Specification and Function in Plants.” Annual Review of Plant Biology 62, no. 1 (June 2, 2011): 461–84. 10.1146/annurev-arplant-042110-103824.

31. Calarco, Joseph P., Filipe Borges, Mark T.A. Donoghue, Frédéric Van Ex, Pauline E. Jullien, Telma Lopes, Rui Gardner, et al. “Reprogramming of DNA Methylation in Pollen Guides Epigenetic Inheritance via Small RNA.” Cell 151, no. 1 (September 2012): 194–205. 10.1016/j.cell.2012.09.001.

32. Borg, Michael, Ranjith K Papareddy, Rodolphe Dombey, Elin Axelsson, Michael D Nodine, David Twell, and Frédéric Berger. “Epigenetic Reprogramming Rewires Transcription during the Alternation of Generations in Arabidopsis.” ELife 10 (January 25, 2021): e61894. 10.7554/eLife.61894.

33. Griffin, Patrick T, Chad E Niederhuth, and Robert J Schmitz. “A Comparative Analysis of 5-Azacytidine-and Zebularine-Induced DNA Demethylation.” G3 Genes|Genomes|Genetics 6, no. 9 (September 1, 2016): 2773–80. 10.1534/g3.116.030262.

34. Liu, Shuo, Yu Bao, Hui Deng, Guanqing Liu, Yangshuo Han, Yuechao Wu, Tao Zhang, and Chen Chen. “The Methylation Inhibitor 5-Aza-2′-Deoxycytidine Induces Genome-Wide Hypomethylation in Rice.” Rice 15, no. 1 (December 2022): 35. 10.1186/s12284-022-00580-6.

35. Zhao, Shuai, Lingling Cheng, Yifei Gao, Baichao Zhang, Xiangdong Zheng, Liang Wang, Pilong Li, Qianwen Sun, and Haitao Li. “Plant HP1 Protein ADCP1 Links Multivalent H3K9 Methylation Readout to Heterochromatin Formation.” Cell Research 29, no. 1 (January 2019): 54–66. 10.1038/s41422-018-0104-9.

36. Lee, Jung-Hoon, Daniel Bollschweiler, Tillman Schäfer, and Robert Huber. “Structural Basis for the Regulation of Nucleosome Recognition and HDAC Activity by Histone Deacetylase Assemblies.” Science Advances 7, no. 2 (January 6, 2021): eabd4413. 10.1126/sciadv.abd4413

37. Wang, Ming, Zhenhui Zhong, Javier Gallego-Bartolomé, Zheng Li, Suhua Feng, Hsuan Yu Kuo, Ryan L. Kan, et al. “A Gene Silencing Screen Uncovers Diverse Tools for Targeted Gene Repression in Arabidopsis.” Nature Plants 9, no. 3 (March 6, 2023): 460–72. 10.1038/s41477-023-01362-8.

38. Andrews, Forest H., Qiong Tong, Kelly D. Sullivan, Evan M. Cornett, Yi Zhang, Muzaffar Ali, JaeWoo Ahn, et al. “Multivalent Chromatin Engagement and Inter-Domain Crosstalk Regulate MORC3 ATPase.” Cell Reports 16, no. 12 (September 2016): 3195–3207. 10.1016/j.celrep.2016.08.050.

39. Li, Sisi, Linda Yen, William A. Pastor, Jonathan B. Johnston, Jiamu Du, Colin J. Shew, Wanlu Liu, et al. “Mouse MORC3 Is a GHKL ATPase That Localizes to H3K4me3 Marked Chromatin.” Proceedings of the National Academy of Sciences 113, no. 35 (August 30, 2016). 10.1073/pnas.1609709113.

40. Zhang, Yi, Brianna J. Klein, Khan L. Cox, Bianca Bertulat, Adam H. Tencer, Michael R. Holden, Gregory M. Wright, et al. “Mechanism for Autoinhibition and Activation of the MORC3 ATPase.” Proceedings of the National Academy of Sciences 116, no. 13 (March 26, 2019): 6111–19. 10.1073/pnas.1819524116.

41. Liu, Yanli, Wolfram Tempel, Qi Zhang, Xiao Liang, Peter Loppnau, Su Qin, and Jinrong Min. “Family-Wide Characterization of Histone Binding Abilities of Human CW Domain-Containing Proteins.” Journal of Biological Chemistry 291, no. 17 (April 2016): 9000–9013. 10.1074/jbc.M116.718973.

42. Hoppmann, Verena, Tage Thorstensen, Per Eugen Kristiansen, Silje Veie Veiseth, Mohummad Aminur Rahman, Kenneth Finne, Reidunn B Aalen, and Rein Aasland. “The CW Domain, a New Histone Recognition Module in Chromatin Proteins: H3K4me Recognition by the CW Domain.” The EMBO Journal 30, no. 10 (May 18, 2011): 1939–52. 10.1038/emboj.2011.108.

43. Young, Matthew D, and Sam Behjati. “SoupX Removes Ambient RNA Contamination from Droplet-Based Single-Cell RNA Sequencing Data.” GigaScience 9, no. 12 (December 26, 2020): giaa151. 10.1093/gigascience/giaa151.

44. Hao, Yuhan, Stephanie Hao, Erica Andersen-Nissen, William M. Mauck, Shiwei Zheng, Andrew Butler, Maddie J. Lee, et al. “Integrated Analysis of Multimodal Single-Cell Data.” Cell 184, no. 13 (June 2021): 3573–3587.e29. 10.1016/j.cell.2021.04.048.

45. McGinnis, Christopher S., Lyndsay M. Murrow, and Zev J. Gartner. “DoubletFinder: Doublet Detection in Single-Cell RNA Sequencing Data Using Artificial Nearest Neighbors.” Cell Systems 8, no. 4 (April 2019): 329–337.e4. 10.1016/j.cels.2019.03.003.

46. Langmead, Ben, and Steven L Salzberg. “Fast Gapped-Read Alignment with Bowtie 2.” Nature Methods 9, no. 4 (April 2012): 357–59. 10.1038/nmeth.1923.

47. Ramírez, Fidel, Devon P Ryan, Björn Grüning, Vivek Bhardwaj, Fabian Kilpert, Andreas S Richter, Steffen Heyne, Friederike Dündar, and Thomas Manke. “DeepTools2: A next Generation Web Server for Deep-Sequencing Data Analysis.” Nucleic Acids Research 44, no. W1 (July 8, 2016): W160–65. 10.1093/nar/gkw257.

48. Zhang, Yong, Tao Liu, Clifford A Meyer, Jérôme Eeckhoute, David S Johnson, Bradley E Bernstein, Chad Nusbaum, et al. “Model-Based Analysis of ChIP-Seq (MACS).” Genome Biology 9, no. 9 (November 2008): R137. 10.1186/gb-2008-9-9-r137.

49. Dobin, Alexander, Carrie A. Davis, Felix Schlesinger, Jorg Drenkow, Chris Zaleski, Sonali Jha, Philippe Batut, Mark Chaisson, and Thomas R. Gingeras. “STAR: Ultrafast Universal RNA-Seq Aligner.” Bioinformatics 29, no. 1 (January 1, 2013): 15–21. 10.1093/bioinformatics/bts635.

50. Anders, Simon, Paul Theodor Pyl, and Wolfgang Huber. “HTSeq—a Python Framework to Work with High-Throughput Sequencing Data.” Bioinformatics 31, no. 2 (January 15, 2015): 166–69. 10.1093/bioinformatics/btu638.

51. Krueger, Felix, and Simon R. Andrews. “Bismark: A Flexible Aligner and Methylation Caller for Bisulfite-Seq Applications.” Bioinformatics 27, no. 11 (June 1, 2011): 1571–72. 10.1093/bioinformatics/btr167.

52. Huang, Xiaosan, Shaoling Zhang, Kongqing Li, Jyothi Thimmapuram, and Shaojun Xie. “ViewBS: A Powerful Toolkit for Visualization of High-Throughput Bisulfite Sequencing Data.” Edited by Jonathan Wren. Bioinformatics 34, no. 4 (February 15, 2018): 708–9. 10.1093/bioinformatics/btx633.

53. Du, Jiamu, Xuehua Zhong, Yana V. Bernatavichute, Hume Stroud, Suhua Feng, Elena Caro, Ajay A. Vashisht, et al. “Dual Binding of Chromomethylase Domains to H3K9me2-Containing Nucleosomes Directs DNA Methylation in Plants.” Cell 151, no. 1 (September 2012): 167–80. 10.1016/j.cell.2012.07.034.

54. Guo, Yixin, Ziwei Xue, Ruihong Yuan, Jingyi Jessica Li, William A. Pastor, and Wanlu Liu. “RAD: A Web Application to Identify Region Associated Differentially Expressed Genes.” Edited by Lenore Cowen. Bioinformatics 37, no. 17 (September 9, 2021): 2741–43. 10.1093/bioinformatics/btab075.

55. Quinlan, Aaron R., and Ira M. Hall. “BEDTools: A Flexible Suite of Utilities for Comparing Genomic Features.” Bioinformatics 26, no. 6 (March 15, 2010): 841–42. 10.1093/bioinformatics/btq033.

56. Sievers, Fabian, Andreas Wilm, David Dineen, Toby J Gibson, Kevin Karplus, Weizhong Li, Rodrigo Lopez, et al. “Fast, Scalable Generation of High-quality Protein Multiple Sequence Alignments Using Clustal Omega.” Molecular Systems Biology 7, no. 1 (January 2011): 539. 10.1038/msb.2011.75.

57. Goujon, M., H. McWilliam, W. Li, F. Valentin, S. Squizzato, J. Paern, and R. Lopez. “A New Bioinformatics Analysis Tools Framework at EMBL-EBI.” Nucleic Acids Research 38, no. Web Server (July 1, 2010): W695–99. 10.1093/nar/gkq313.

58. McWilliam, Hamish, Weizhong Li, Mahmut Uludag, Silvano Squizzato, Young Mi Park, Nicola Buso, Andrew Peter Cowley, and Rodrigo Lopez. “Analysis Tool Web Services from the EMBL-EBI.” Nucleic Acids Research 41, no. W1 (July 1, 2013): W597–600. 10.1093/nar/gkt376.

59. Letunic, Ivica, and Peer Bork. “Interactive Tree Of Life (ITOL) v5: An Online Tool for Phylogenetic Tree Display and Annotation.” Nucleic Acids Research 49, no. W1 (July 2, 2021): W293–96. 10.1093/nar/gkab301.

60. Yan, Liuhua, Shaowei Wei, Yaorong Wu, Ruolan Hu, Hongju Li, Weicai Yang, and Qi Xie. “High-Efficiency Genome Editing in Arabidopsis Using YAO Promoter-Driven CRISPR/Cas9 System.” Molecular Plant 8, no. 12 (December 2015): 1820–23. 10.1016/j.molp.2015.10.004.

