## Supplementart Figures for "MBD2 couples DNA methylation to Transposable Elements silencing during male gametogenesis"

Supplementary Fig. 1-6

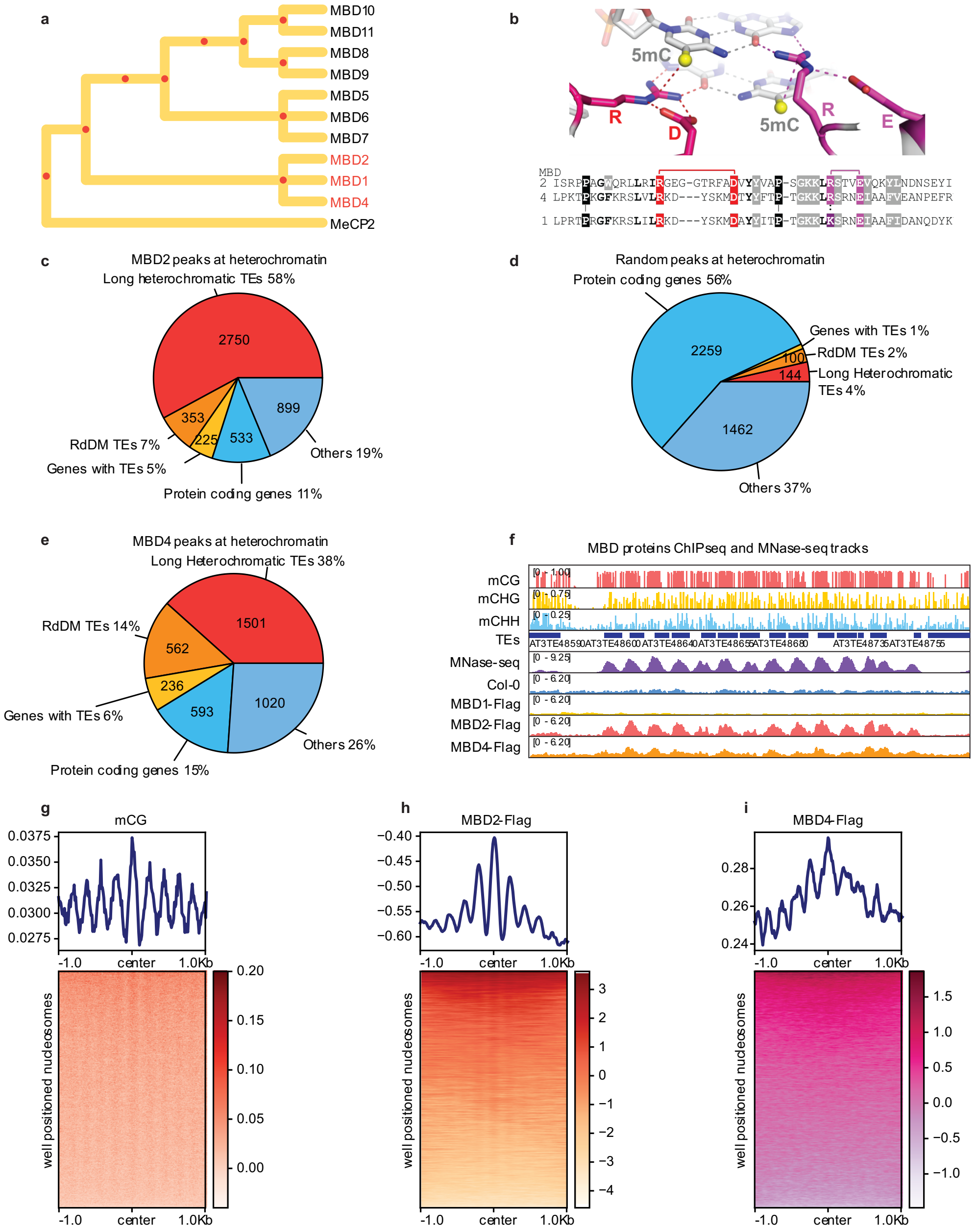


**Supplementary Fig. 1 | MBD2 and MBD4 are novel methyl readers. a.** Phylogenetic tree of the MBD proteins, generated using the conserved sequences within the MBD domains by Clusteral Omega (Table S2). MeCP2 is used as the outgroup. Red dots indicate the hypothetical common ancestors. **b.** Adapted from Figure S3 of Ichino, et al[8]. Protein 3D modeling of the interactions between the MBD domain and the methylated cytosines. MBD domain sequences are shown below, red and purple highlighting the critical amino acids for the interactions. **c-e.** Proportions of heterochromatic peaks called from **c.** MBD2 ChIP-seq, **d.** random shuffling, and **e.** MBD4 ChIP-seq that overlap with long heterochromatic TEs, RdDM-associated TEs, genes with TEs, protein-coding genes, and others. **f.** Screenshot of the ChIP-seq tracks of MNase-seq, MBD1, MBD2, and MBD4 with the methylation level at the representative TE sites with well positioned nucleosomes. **g-h.** Metaplots and heatmaps showing **g.** CG methylation, **h.** MBD2 ChIP-seq, **i.** MBD4 ChIP-seq, and **j.** CMT3 ChIP-seq (data from Du et al.[53]) at well positioned nucleosomes.

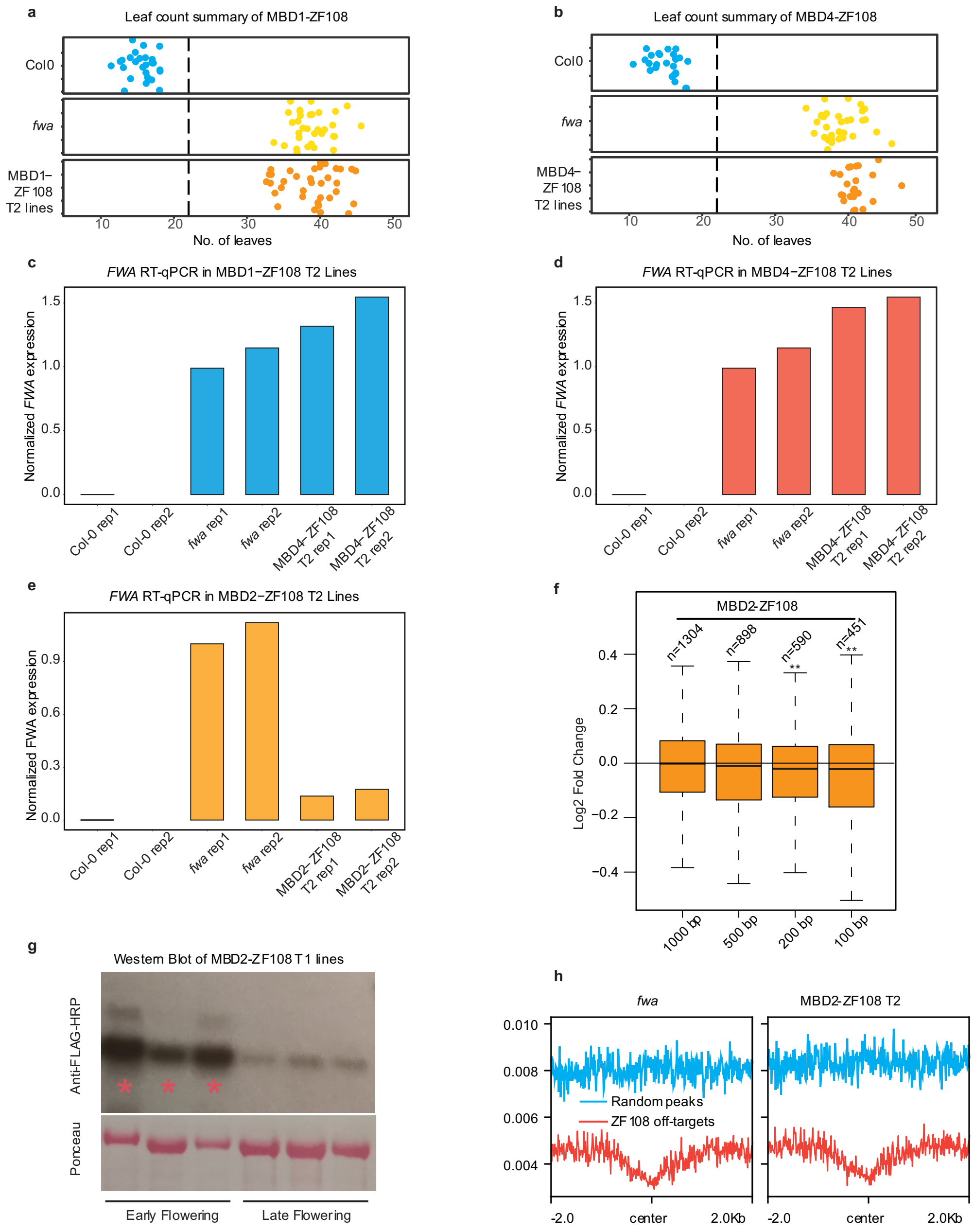


**Supplementary Fig. 2 | MBD2 but not MBD1 and MBD4 silences FWA exogenously. a-b.** Flowering time of Col-0, *fwa*, and the representative T2 lines of **a.** MBD1-ZF108 and **b.** MBD4-ZF108 as measured by the number of leaves. **c-e.** qRT-PCR analysis showing the relative mRNA level of *FWA* in Col-0, *fwa*, and the representative T2 lines of **c.** MBD1-ZF108, **d.** MBD4-ZF108, and **e.** MBD2-ZF108. **f.** Log2 fold change of the DEGs close to ZF108 off-target sites from MBD2-ZF108 leaf RNA-seq. P value is calculated by student t-test, P<0.01: ** **g.** Western blot of the representative early flowering and late flowering T1 lines of MBD2-ZF108, which were used for flowering time measurement in Fig.1e. Pink asterisk indicates early flowering T2 plants. **h.** CG methylation level at the ZF108 off-target sites and random sites of *fwa* and the representative T2 line of MBD2-ZF108.

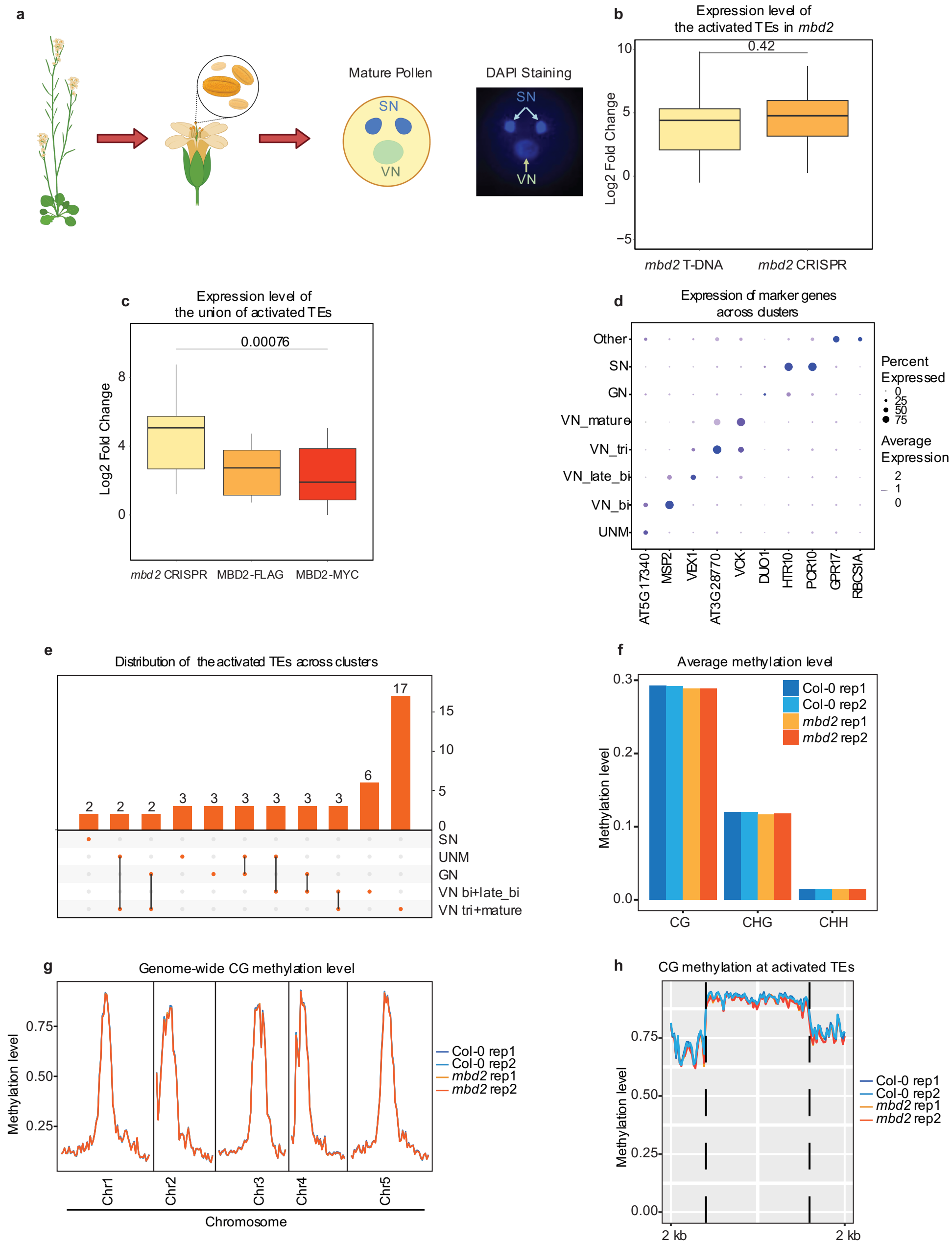


**Supplementary Fig. 3 | MBD2 works downstream of DNA methylation to prevent TE activation during male gametogenesis. a.** Graphic illustration showing the morphology and DAPI staining of wild-type mature pollen. **b.** Log2 fold change of the activated TEs in *mbd2* T-DNA and CRISPR mutants. **c.** Dot plot showing the cluster specificity using the expression of known markers. The dot size represents the percentage of cells in which the gene was detected. The dot color corresponds to the scaled average expression. **d.** Upset plot showing the distribution of activated TEs in the *mbd2* mutant across different clusters. **e-f.** Genome-wide CG, CHG, and CHH methylation levels of Col-0 and *mbd2* mature pollen. **g.** CG methylation level at the mbd2 activated TEs of Col-0 and *mbd2*.

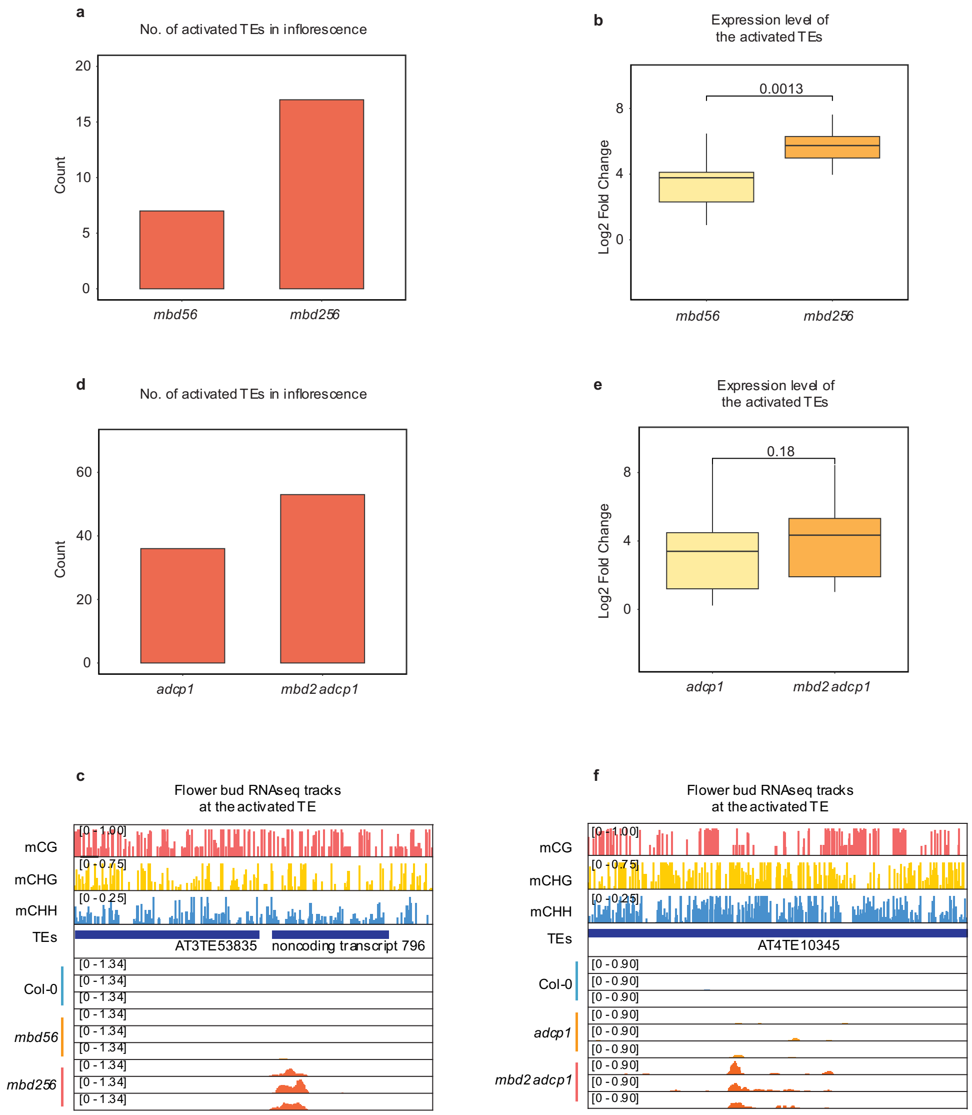


**Supplementary Fig. 4 | MBD2 plays a repressive role at TE in sensitized backgrounds. a.** Count of the activated TEs of *mbd56* and *mbd256* from inflorescence RNA-seq. **b.** Log2 fold change of the activated TEs of *mbd56* and *mbd256* from inflorescence RNA-seq. **c.** Screenshot of the inflorescence RNA-seq tracks of Col-0, *mbd56*, and *mbd256* with the methylation level at the representative TE and the DNA methylated transcript. **d.** Count of the activated TEs of *adcp1* and *mbd2 adcp1* from inflorescence RNA-seq. **e.** Log2 fold change of the activated TEs of *adcp1* and *mbd2 adcp1* from inflorescence RNA-seq. **f.** Screenshot of the inflorescence RNA-seq tracks of Col-0, *adcp1*, and *mbd2 adcp1* with the methylation level at the representative TE.

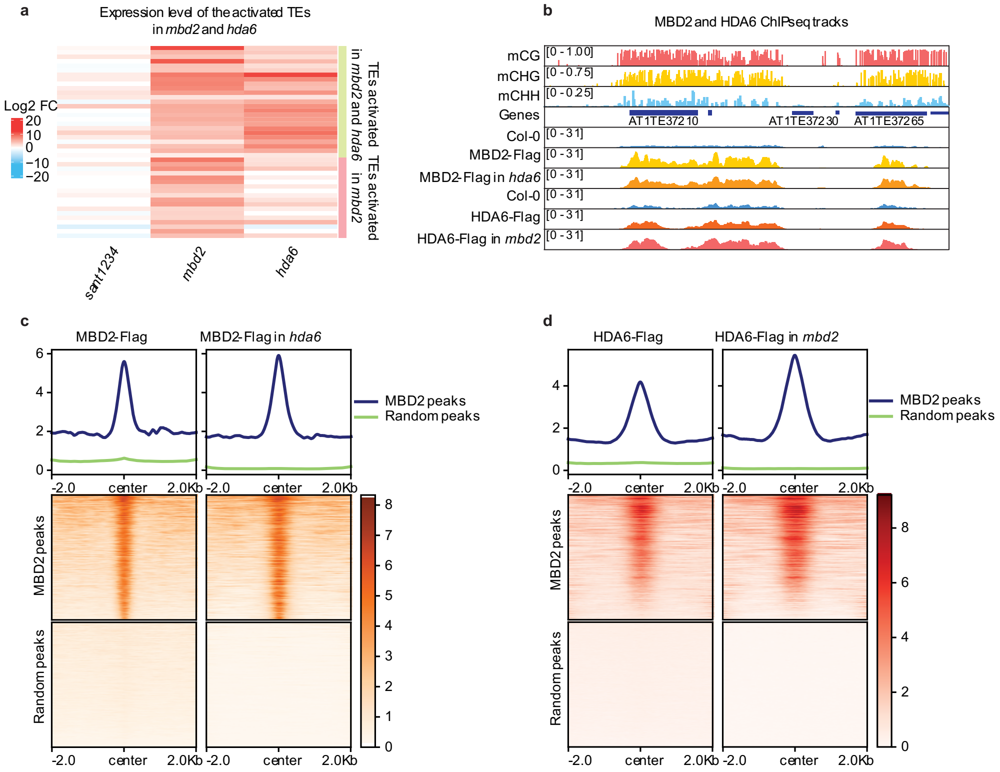


**Supplementary Fig. 5 | MBD2 mediated silencing is independent of HDA6 and SANT family proteins. a.** Heatmaps showing the log2 fold change of the activated TEs in *sant1234*, *mbd2*, and *hda6* from mature pollen RNA-seq. **b.** Screenshot of the ChIP-seq tracks of MBD2, MBD2 in *hda6*, HDA6, and HDA6 in *mbd2* with methylation level at the representative TEs. **c-f.** Metaplots and heatmaps showing the ChIP-seq signal of **c.** MBD2, MBD2 in *hda6*, **d.** HDA6, and HDA6 in *mbd2* at the DNA methylated MBD2 peaks shared with HDA6 and random peaks.

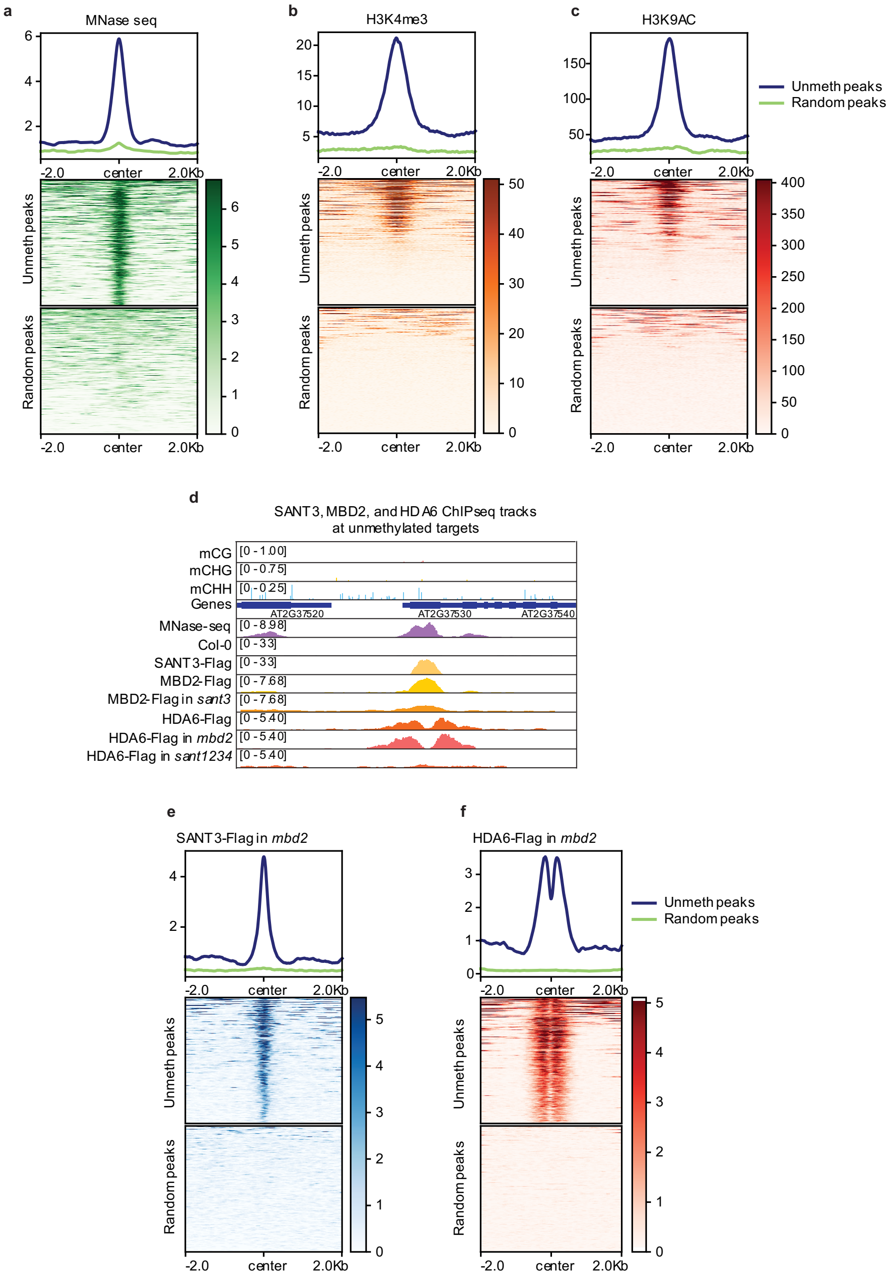


**Supplementary Fig. 6 | MBD2 forms a complex with HDA6 and SANT family proteins at +1 nucleosomes of unmethylated genes. a-c.** Metaplots and heatmaps showing **a.** MNase-seq, **b.** H3K4me3, and **c.** H3K9AC enrichment at the unmethylated peaks and random peaks. **d.** Screenshot of the ChIP-seq tracks of the indicated samples with methylation level at the representative unmethylated genes. **e-f.** Metaplots and heatmaps showing the ChIP-seq signal of **e.** SANT3 in *mbd2*, and **f.** HDA6 in *mbd2* at the unmethylated peaks and random peaks.

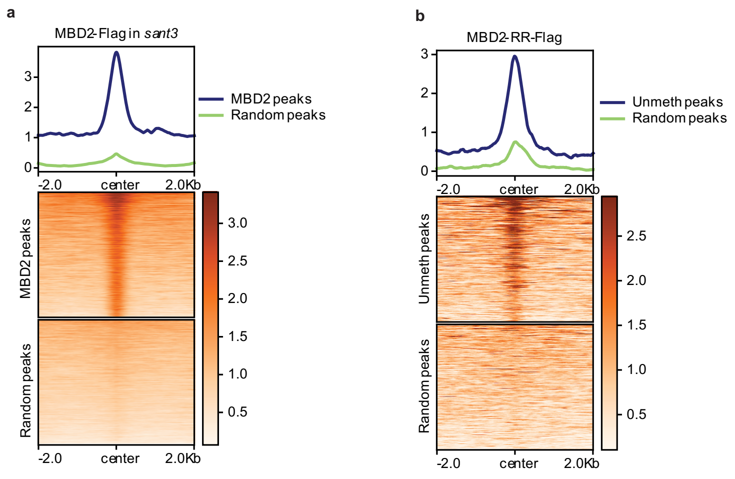


**Supplementary Fig. 7 | Different mechanisms mediate MBD2 localization to unmethylated and methylated regions.** Metaplots and heatmaps showing the ChIP-seq signal of **a.** MBD2 in *sant3* at the DNA methylated MBD2 peaks shared with HDA6 and random peaks, and **b**. MBD2-RR at unmethylated peaks shared among MBD2, SANT3, and HDA6, and random peaks.

**a**

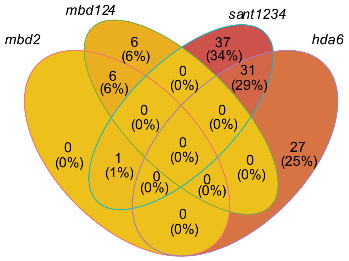


**Supplementary Fig. 8 | MBD1, MBD2, and MBD4 are dispensable for SANT and HDA6 involved gene regulation. a.** Venn diagram showing the overlap of activated unmethylated DEGs among *mbd2*, *mbd124*, *sant1234*, and *hda6*.
